# A small-molecule inhibitor of the BRCA2-RAD51 interaction modulates RAD51 assembly and potentiates DNA damage-induced cell death

**DOI:** 10.1101/2020.07.15.200709

**Authors:** Duncan E. Scott, Nicola J. Francis-Newton, May E. Marsh, Anthony G. Coyne, Gerhard Fischer, Tommaso Moschetti, Andrew R. Bayly, Timothy D. Sharpe, Kalina T. Haas, Lorraine Barber, Chiara R. Valenzano, Rajavel Srinivasan, David J. Huggins, Matthias Ehebauer, Alessandro Esposito, Luca Pellegrini, Trevor Perrior, Grahame McKenzie, Tom L. Blundell, Marko Hyvönen, John Skidmore, Ashok R. Venkitaraman, Chris Abell

**Affiliations:** Department of Chemistry, Lensfield Road, University of Cambridge, Cambridge, CB2 1EW, UK; Medical Research Council Cancer Unit, University of Cambridge, Hutchison/MRC Research Centre, Hills Road, Cambridge CB2 0XZ, UK; Department of Biochemistry, University of Cambridge, 80 Tennis Court Road, Cambridge CB2 1GA, UK; The ALBORADA Drug Discovery Institute, Island Research Building, Hills Road, University of Cambridge, Cambridge, UK, CB2 0AH; Paul Scherrer Institut, Villigen PSI, Switzerland; Boehringer Ingelheim RCV, Doktor-Boehringer-Gasse 5-11, 1121 Vienna, Austria; Illumina Cambridge Ltd, Illumina Centre, Granta Park, Great Abington, Cambridge, CB21 6GP, UK; Vertex, 86-88 Jubilee Avenue, Milton Park, Abingdon, Oxfordshire OX14 4RW; Biophysics Facility, Biozentrum, University of Basel, Basel, Switzerland; Astex Pharmaceuticals, 436 Cambridge Science Park, Milton Road, Cambridge, CB4 0QA, U.K.; School of Pharmaceutical Science and Technology, Health Science Platform, Tianjin University, 92 Weijin Road, Nankai District, Tianjin 300072; People’s Republic of China; Department of Physiology and Biophysics, Weill Cornell Medical College, USA; Ipsen Bioinnovation, Oxford; Domainex, Chesterford Research Park, Little Chesterford, Saffron Walden, Essex, CB10 1XL, UK; PhoreMost Ltd., Babraham Research Campus, Cambridge CB22 3AT, UK

**Keywords:** RAD51, homologous recombination, BRCA2, DNA repair, structure-guided drug discovery, protein-protein interaction inhibition

## Abstract

BRCA2 controls RAD51 recombinase during homologous DNA recombination (HDR) through eight evolutionarily-conserved BRC repeats, which individually engage RAD51 via the motif Phe-x-x-Ala. Using structure-guided molecular design, templated on a monomeric thermostable chimera between human RAD51 and archaeal RadA, we identify CAM833, a 529 Da orthosteric inhibitor of RAD51:BRC with a K_d_ of 366 nM. The quinoline of CAM833 occupies a hotspot, the Phe-binding pocket on RAD51 and the methyl of the substituted α-methylbenzyl group occupies the Ala-binding pocket. In cells, CAM833 diminishes formation of damage-induced RAD51 nuclear foci; inhibits RAD51 molecular clustering, suppressing extended RAD51 filament assembly; potentiates cytotoxicity by ionising radiation, augmenting *4N* cell-cycle arrest and apoptotic cell death and works with poly-ADP ribose polymerase (PARP)1 inhibitors to suppress growth in BRCA2-wildtype cells. Thus, chemical inhibition of the protein-protein interaction between BRCA2 and RAD51 disrupts HDR and potentiates DNA damage-induced cell death, with implications for cancer therapy.

## INTRODUCTION

The tumour suppressor protein, BRCA2, is essential for error-free repair of DNA double-stranded breaks (DSBs) by homologous DNA recombination (HDR) in human cells (Venkitaraman, 2014). BRCA2 acts during HDR to control the recombination enzyme, RAD51, a eukaryal protein evolutionarily conserved as RecA in eubacteria, and RADA in archaea (West, 2003). RAD51 executes the DNA strand exchange reactions that lead to HDR by assembling, in a sequential and highly regulated manner, as helical nucleoprotein filaments on single-stranded (ss) or double-stranded (ds) DNA substrates. The presynaptic RAD51 filament on ssDNA mediates strand invasion and homologous pairing with a duplex DNA template to execute strand exchange, the core biochemical event necessary for HDR.

Human BRCA2 contains two distinct regions that bind directly to RAD51. First, BRCA2 contains eight BRC repeats, evolutionarily-conserved motifs of 26 residues each, whose sequence and spacing within an ∼1,100 residue segment encoded by *BRCA2* exon 11 is conserved amongst mammalian orthologues (Bignell et al., 1997). Each of the eight human BRC repeats exhibits a varying affinity for RAD51 *in vitro* (Wong et al., 1997). Second, the carboxyl (C-)terminus of BRCA2 contains a RAD51-binding region spanning ∼90 residues, which is distinct in sequence from the BRC repeats (Davies and Pellegrini, 2007; Esashi et al., 2007).

The interactions between BRCA2 and RAD51 control key steps essential for HDR. The BRC repeat-RAD51 interaction differentially regulates RAD51 assembly on DNA substrates *in vitro*, promoting assembly of the RAD51-ssDNA filament, whilst concurrently inhibiting the RAD51-dsDNA interaction (Carreira et al., 2009; Shivji et al., 2009). These opposing activities of the BRC repeats ensure that RAD51 assembly on its DNA substrates occurs in the correct order to promote strand exchange. Moreover, the C-terminal RAD51-binding region of BRCA2 stabilizes oligomeric assemblies of RAD51 *in vitro* in biochemical assays using purified proteins (Davies and Pellegrini, 2007; Esashi et al., 2007), and is required for the elongation of RAD51 filaments at cellular sites of DNA damage visualized by single-molecule localization microscopy (Haas et al., 2018).

Of the eight BRC repeats in human BRCA2, BRC4 exhibits the highest affinity for RAD51 (Carreira and Kowalczykowski, 2011; Cole et al., 2017; Wong et al., 1997). The crystallographic structure of a complex between a BRC4 peptide and the core catalytic domain of RAD51 shows that the BRC4 sequence FHTA (human BRCA2 residues 1524-1527) engages with hydrophobic pockets on the RAD51 surface that accommodate the Phe and Ala residues (Pellegrini et al., 2002). An analogous FxxA motif in the RAD51 protein mediates oligomerization in the absence of DNA (Brouwer et al., 2018; Conway et al., 2004; Shin et al., 2003), and has recently been shown using electron cryo-micoscopy to form the inter-subunit interface in functionally relevant DNA-bound assemblies of RAD51 (Short et al., 2016; Xu et al., 2017). *In vitro*, BRC4 peptides promote the strand selectivity of RAD51-DNA interactions at sub-stoichiometric concentrations relative to RAD51 (Carreira et al., 2009; Shivji et al., 2009). However, BRC4 peptides disrupt RAD51 oligomerization *in vitro* (Davies et al., 2001), and when overexpressed in cells, can inhibit the recruitment of RAD51 into DNA damage-induced foci by blocking the RAD51-RAD51 interaction (Chen et al., 1999a).

The central importance of the BRC repeat-RAD51 interaction to HDR has prompted the identification of small-molecule and peptidic inhibitors that might have therapeutic value for cancer treatment. Most reported inhibitors target the interaction between RAD51 and DNA (Budke et al., 2012a, 2012b; Huang and Mazin, 2014; Huang et al., 2011; Ishida et al., 2009; Normand et al., 2014; Takaku et al., 2011). Recently described cell penetrating antibodies also operate through a similar mechanism (Pastushok et al., 2019; Turchick et al., 2017, 2019). Inhibitors that suppress the D-loop activity of RAD51 have also been reported (Budke et al., 2019; Lv et al., 2016), although several optimized versions also exhibit DNA-intercalating activity (Budke et al., 2019). On the other hand, reports of small molecules and peptides have been published that claim to disrupt the interaction between RAD51 and the BRC repeats, or between RAD51 multimers (Bagnolini et al., 2020; Falchi et al., 2017; Nomme et al., 2010; Roberti et al., 2019; Trenner et al., 2018; Vydyam et al., 2019; Zhu et al., 2013, 2015; Ward et al. 2017). However, the lack of specific structural information concerning the interaction of these inhibitors with RAD51 has impeded the precise exploration of structure-activity relationships, and the efficient development of more potent compounds.

Here, we report the discovery, using a structure-led fragment-based approach, of CAM833, a potent chemical inhibitor of the RAD51-BRC repeat interaction and RAD51 oligomerization. We show using X-ray crystallography that CAM833 engages the Phe-and Ala-binding pockets on RAD51 to block its interaction with BRC repeats. We confirm that CAM833 potentiates cellular sensitivity to DNA damage induced by ionizing radiation, and suppresses the assembly of RAD51 into damage-induced filaments, as visualized by single-molecule localization microscopy. Our findings provide a well characterized chemical tool compound to dissect biochemical events during HDR, and a potential lead for the development of new cancer therapeutics.

## RESULTS

### A monomeric thermostable chimera of human RAD51 and archaeal RADA recapitulates structural features of the human RAD51-BRC interaction

Structure-based approaches to identify modulators of the BRCA2–RAD51 interaction have been impeded by the lack of a monomeric unliganded form of *Hs*RAD51. We have previously described the development of molecular surrogate systems for RAD51 based on an archaeal ortholog, RadA from *Pyrococcus furiosus* (Moschetti et al., 2016). In brief, we were able to produce the C-terminal ATPase domain of RadA as a stable monomeric protein, and by careful mutagenesis, to convert the surface of the protein to resemble human RAD51, with the ability to bind the BRCA2 BRC4 repeat with high affinity. Of note, we used the previously described constructs HumRadA2 for initial biophysics work and HumRadA22F for crystallography (Moschetti et al., 2016). In parallel, we also generated a chimeric RAD51 (ChimRAD51) that fuses the central part of the human RAD51 ATPase domain with two flanking sequences from archaeal RadA and used this in our primary screening assay and for biophysical screening. The binding of ChimRAD51 to the BRC4 peptide was characterized using a fluorescence polarization (FP) assay and by isothermal titration calorimetry (ITC), yielding comparable K_d_ values of 4 and 6 nM, respectively validating the use of this protein for subsequent ligand affinity measurements (Moschetti et al., 2016). These surrogates provide robust platforms for structure-guided lead discovery via fragment screening, the biophysical characterization and validation of inhibitors, and for X-ray crystallography.

The three-dimensional structure of the C-terminal ATPase domain of RAD51 in complex with a BRC4 peptide has been determined by X-ray crystallography (Pellegrini et al., 2002). This structure shows that the BRC repeat binds in a bi-dentate fashion in which BRC4, via its FxxA motif, engages with a self-association site on RAD51, and then wraps around the protein to interact through a less-conserved LFDE motif with a second site on the RAD51 surface (Figure 1A). Cryo-EM structures of RAD51 filaments bound to DNA (Short et al., 2016; Xu et al., 2017) confirm that in self-association, the FxxA motif of one RAD51 interacts similarly with the two small “Phe” and “Ala” pockets on an adjacent protein unit, with the C-terminal segment of the oligomerization epitope binding to a hydrophobic groove in the opposite direction to that where the LFDE epitope of BRC4 binds. Earlier work has compared the relative affinities of the different human BRC repeats for RAD51 (*eg*. (Wong et al., 1997)), and has demonstrated that both the FxxA and LFDE motifs in multiple BRC repeats contribute to both permissive and inhibitory interactions with RAD51 (Rajendra and Venkitaraman, 2010). In order to determine which of these two motifs might be most appropriate to develop inhibitors against, we measured the affinities of two peptides corresponding to N-and C-terminal epitopes of BRC4 using our FP assay. The N-terminal “FxxA” half of the BRC4 repeat showed clear binding to RAD51 and competition of full-length BRC4 repeat with a K_d_ of 36 μM. This compares favorably with our previous analysis of the affinities of tetra-peptides derived from the BRC4 FxxA epitope (which has the sequence FHTA) which bound to humanized RadA with 200-300 μM affinity (Scott et al., 2016). The C-terminal half of BRC4 (LFDE peptide) showed very weak, if any, inhibition of BRC4 binding, at up to 1 mM concentration (Figure 1C), suggesting that the LFDE motif makes a minimal contribution on its own to this interaction, even though its mutation in the context of the entire BRC4 peptide can reduce RAD51 binding (Rajendra and Venkitaraman, 2010). We also tested the ability of RAD51 to bind its own oligomerization peptide (OP) epitope and determined a K_d_ of 18 μM for this interaction, demonstrating how additional binding energy can be derived from the interactions which the C-terminal part of the oligomerization peptide makes (Figure 1C).

**Figure 1.**
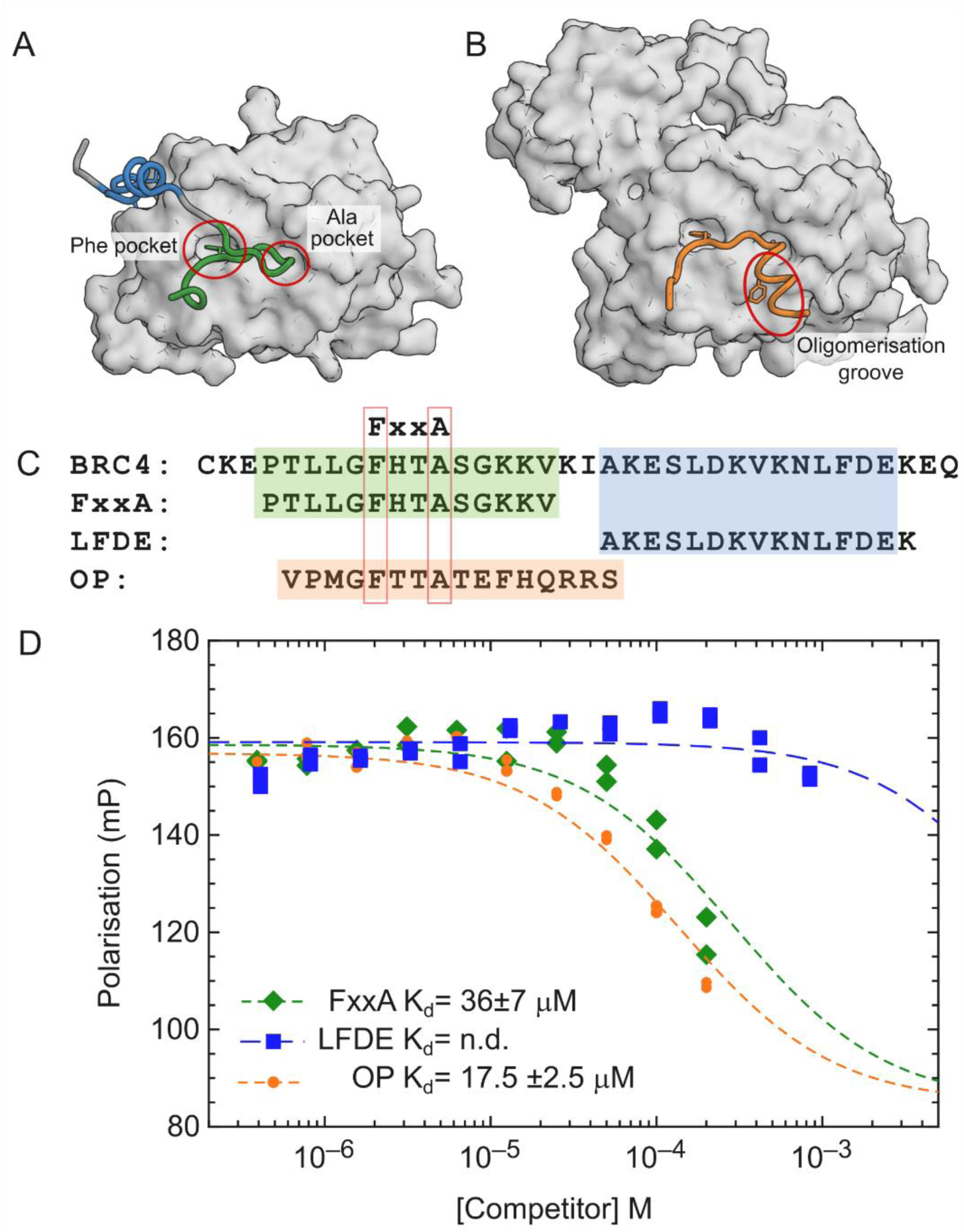
RAD51 interaction with BRC4. A. Structure of RAD51 ATPase domain (surface) with BRC4 repeat of BRCA2 with FxxA binding motif coloured green and the LFDE-motif in blue (PDB code 1n0w). B. Structure of oligomeric RAD51 with oligomerization epitope (orange) of one protomer binding the next molecule in the filament (surface) (PDB 5nlw). C. Sequences of BRC4 repeat, and its FxxA and LFDE epitopes containing half peptides and the isolated RAD51 oligomerization peptide (OP). D. Competitive FP assay with labelled BRC4 repeat as probe which shows competitive binding to ChimRAD51 protein with the two BRC4 half-peptides (FxxA and LFDE, green and blue) and RAD51 oligomerization peptide (OP, orange). The dissociation constants measured for the FxxA half-peptide and for the oligomerization peptides were 36 ± 7 μM and 18 ± 3 μM, respectively.

### The design and development of CAM833, a small molecule inhibitor of the interaction between BRCA2 and RAD51

Using the surrogate RAD51 systems described above and a combination of fragment-based drug discovery (Blundell et al., 2002; Coyne et al., 2010) and structure-guided drug design, we have optimized fragment hit molecules to generate high-affinity inhibitors of the RAD51–BRC-repeat interaction with a clearly defined orthosteric inhibition mechanism (Figure 2).

**Figure 2.**
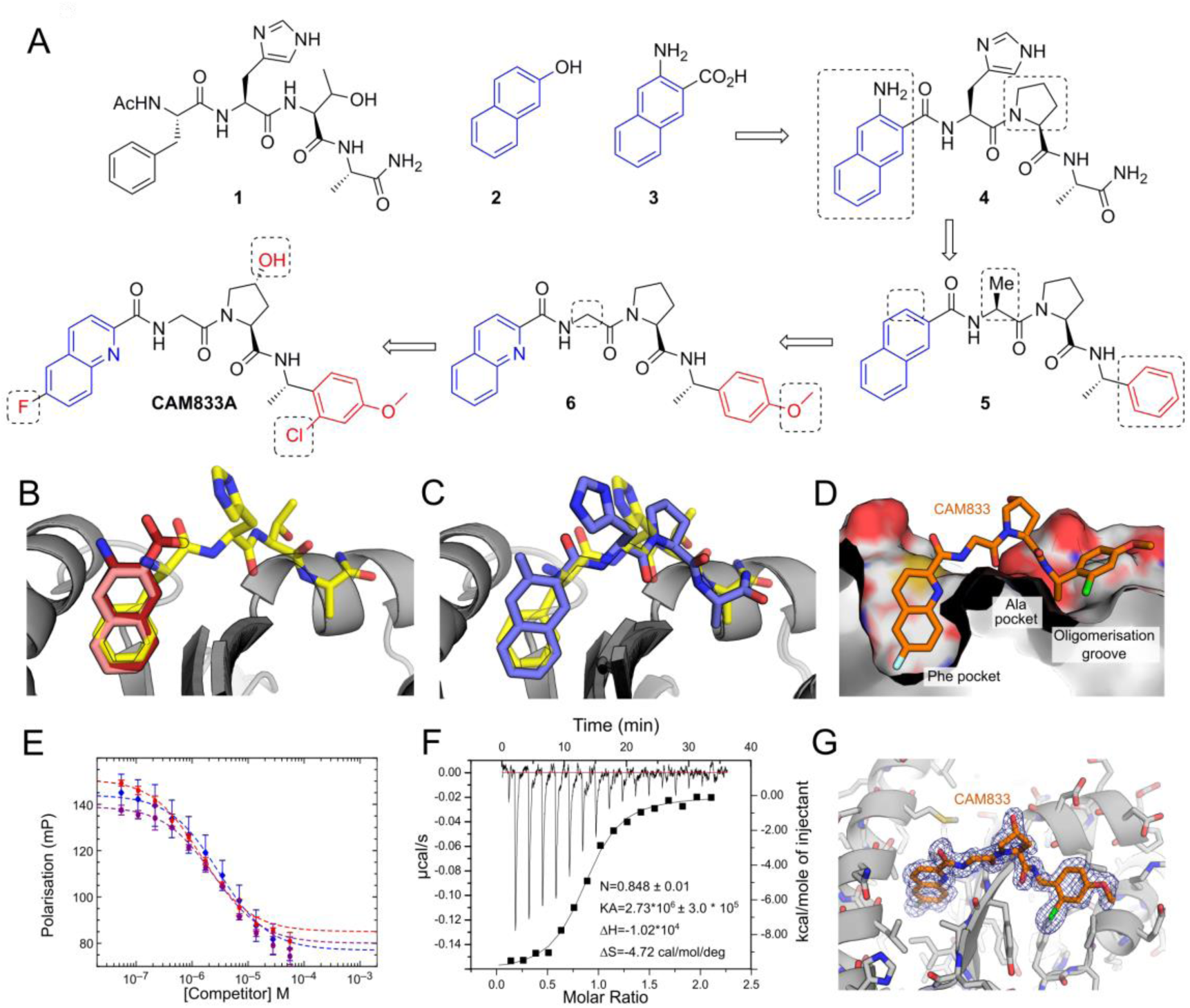
Development of CAM833. (A) Merging of 3-amino-2-naphthoic acid (**3**) with FHPA tetrapeptide (**1**) to yield **4**. Trimming of the naphthyl and histidine group and replacement of terminal amide with phenyl group yields **5**. Increase of polarity by replacing naphthyl with quinoline and adding methoxy group the phenyl ring results in **6**. Further optimization leads to CAM833. (B) Overlaid crystal structures of HumRadA1 in complex with 2-naphthol (**2**, PDB: 4B32, pink), 3-amino-2-naphthoic acid (**3**, PDB:6TV3, dark red) and FHTA tetrapeptide (**1**, PDB: 4B3B, yellow). (C) Structure of **4** (PDB: 6TWE, deep purple) in complex with HumRadA1 overlaid with FHTA peptide (PDB: 4B3B, yellow). (D) Structure of CAM833 (orange, PDB: 6TW9) in complex with HumRadA22F. Side view of CAM833 complex with HumRadA22F showing partially cut surface of the protein and interaction of the fluoroquinoline ring with the Phe-pocket and the chloro-phenyl group binding into the oligomerization groove. (E) Competition of BRC4 peptide binding to ChimRAD51 using FP assay with CAM833. Three independent measurements (triplicate technical repeats) of the same binding are shown in three different colours. (F) Isothermal titration calorimetric measurement of direct binding of CAM833 to ChimRAD51. The baseline corrected thermogram is shown above with X-and Y-axes above and left of the graph. The solid squares depict integrated heats for each titration point and solid line the fit to single-site binding mode with corresponding X-and Y-axes below and to left of the graph. (F) Refined 2F_o_F_c_ electron density is shown for the ligand, contoured at 1σ.

Initially, we carried out a biophysical fragment-screen against a previously-described humanized version of *Pf*RadA HumRadA2 (also known as MAYSAM; (Moschetti et al., 2016; Scott et al., 2013)) leading to the discovery of a range of bicyclic aromatic and heteroaromatic fragment hits, binding exclusively into the Phe pocket at the FxxA site of RAD51 (Scott et al., 2013). Investigation of the structure-activity relationships (SAR) around these hits showed that naphthyl derivatives, particularly when substituted with polar groups, were able to bind to the Phe pocket with reasonable activity and ligand efficiency. For example, 2-hydroxynaphthalene (**2**) bound to the HumRadA2 protein with a K_d_ of 460 μM as measured by ITC (Scott et al., 2013), whereas 3-amino-2-naphthoic acid (**3**) (Figure 2A) bound with a Kd of 1.36 mM (Supplementary figure S1). Crystallographic analysis of these fragments shows that the naphthyl rings bind in the same orientation as the aromatic side chain of phenylalanine in the FxxA motif of oligomerization peptide or BRC repeats (Figure 2B) (Scott et al., 2016).

In parallel, we explored the SAR of a series of *N*-acetylated tetrapeptides based on the FxxA epitope of BRC4 (Scott et al., 2016). This work established that the Ac-FHTA-NH_2_ tetrapeptide (**1**) binds to HumRadA2 with a K_d_ of 280 μM as determined by ITC.

Based on an overlay of the X-ray crystal structures of **1** and the naphthyl fragments 2-hydroxynaphthalene (**2**) and 3-amino-2-naphthoic acid (**3**) (figure 2B) we designed compound **4** in which the Phe of FHTA has been replaced by a rigid naphthyl-based amino acid, designed to more completely fill this pocket, and the threonine has been replaced by a proline in order to restrict the conformation of the peptide. The latter modification is known to provide a modest potency increase from the tetrapeptide structure activity relationship studies (Scott et al., 2016), with the benefit of removing two H-bond donors from the structure, a change likely to be associated with an increase in cell permeability. Gratifyingly, the merged compound **4** was found to have improved K_d_ of 3 μM against HumRadA2 as determined by ITC (Supplementary figure S1), a considerable increase in potency compared to the native peptide. We determined an X-ray crystal structure of **4** bound to the HumRadA2 protein and this was found to interact in the predicted fashion, with a modest distortion of the peptide backbone in order to accommodate the more rigid left-hand-side (orientation as in Figure 2) (Figure 2C).

Recognizing that the peptidic nature of **4** was likely to lead to poor pharmacokinetics and low permeability (clogP as calculated with ChemDraw 16 -0.96 and tPSA 172 Å^2^), we sought to reduce the size and polarity of our compounds whilst introducing groups capable of forming additional interactions with the protein surface. This led to the design of compound **5** in which three polar elements judged unnecessary were removed: firstly, the His residue which makes no key interactions in the tetrapeptide-protein crystal structure was cut back to an alanine; secondly, we removed the amino group from the terminal naphthoate unit. Finally, the terminal Ala amide was replaced with an α-methylbenzylamino group that maintains the methyl group important for binding into the Ala pocket whilst replacing the terminal-amide with a lipophilic phenyl ring, inspired by the relatively non-polar nature of this region of the protein surface. Overall, compound **5** has only two intact amino acids and greatly reduced polarity (clogP 4.08, tPSA = 78.5). Compound **5** has a K_d_ of 220 nM vs HumRadA2 by ITC (Supplementary figure S2) – a 10-fold potency increase. By this stage we had developed the more thoroughly humanized form of the protein ChimRAD51 which was subsequently used for our primary FP screening assays (Moschetti et al., 2016). We determined the K_d_ of **5** against ChimRAD51 to be 1.9 μM by ITC and 27 μM (n=20) as measured by FP. The reduced level of potency against this more humanized system was mirrored in data with the original naphth-2-ol fragment (**2**) which we found to have a K_d_ of 3.3 mM for ChimRAD51 versus 460 μM for HumRadA2.

Compound **5** was too insoluble in aqueous media to profile in cell-based assays. Accordingly, we made modifications designed to increase polarity, whilst avoiding the introduction of further hydrogen-bond donors likely to reduce permeability. We replaced the naphthyl ring with a quinoline, converted the Ala residue into a Gly and introduced a 4-methoxy substituent on the right-hand phenyl ring, leading to **6**. Compound **6** has a clogP of 2.76 and an improved FP K_d_ of 8.0 μM (n=22) against ChimRAD51. Two independent X-ray structures of **6** demonstrated that this compound was still bound to the FxxA site with the quinoline accessing the Phe pocket in a similar orientation to the naphthyl in compound **4** albeit with a shifted binding mode discussed in more detail below (Supplementary figure S3).

More detailed exploration of the SAR around **6** led to the discovery of **CAM833** with a 6-fluoro substituent on the quinoline and a 2-chloro group on the phenyl leading to a further increase in affinity. CAM833 has a K_d_ against the ChimRAD51 protein of 350 nM (n=8) as measured by FP (Figure 2E) and 366 nM by ITC (Figure 2F). The lipophilicity associated with these groups was balanced by the introduction of a *trans*-4-hydroxyl substituent on the proline ring serving to maintain solubility (clogP of CAM833 is 2.73, and tPSA 120 Å^2^).

As a biochemical test of CAM833, we evaluated its ability to disrupt full-length RAD51 oligomers. Using dynamic light scattering, we observe a shift of the average particle size from ∼40 nm for oligomeric RAD51 to ∼5 nm particles (corresponding closely to the size of a RAD51 monomer) in the presence of excess of CAM833 (Supplementary figure S4).

The X-ray crystal structures of **6** and CAM833 bound to HumRadA22F (the fully humanized RadA surrogate used for crystallography (Moschetti et al., 2016); Figure 2D, G, Figure S3) revealed an altered binding-mode compared to the lead compound **4** (Figure 2C). In this new binding mode, a shift of the backbone of CAM833 allows the NH of the right-hand benzyl amide to form a hydrogen bond to Val200_189_ (subscript number refers to the equivalent human RAD51 residue, which differs from the surrogate protein residue numbering) via a bridging water-molecule rather than directly to the protein backbone (Supplementary figure S5). We attribute this to the truncation of the His residue back to a Gly, a change that can be tracked in the X-ray structures of intermediates from the optimization bound to HumRadA22F (data not shown).

Overall, examination of the structures reveals the source of the potency increases between the tetrapeptide **1** and CAM833. The phenyl ring of CAM833 sits flat on the protein surface with the *ortho-* chlorine atom sitting in a groove leading from this surface with both making beneficial hydrophobic interactions. The quinoline more completely fills the Phe pocket and the 6-fluoro substituent binds into a hydrophobic sub-pocket which has opened up due to minor movements in the residues lining the pocket (Figure 2D and 2G). We determined selectivity and developability data for CAM833 in order to support its use as a validated chemical probe for the RAD51-BRCA-2 protein-protein interaction. Briefly, CAM833 is metabolically stable, does not significantly inhibit CYP450 enzymes, shows moderate solubility and permeability and has no significant off-target interactions when screened at 10 μM in the Cerep ExpresSPanel and has mouse pharmacokinetic data suitable for *in vivo* investigation (Supplementary Table S1).

### CAM833 causes a concentration-dependent decrease in RAD51 foci accompanied by increased DNA damage

The assembly of RAD51 into microscopic foci at cellular sites of DNA damage is competitively inhibited by the over-expression of BRC repeat peptides (Chen et al., 1999a). Indeed, structural studies using X-ray crystallography (Brouwer et al., 2018; Pellegrini et al., 2002; Shin et al., 2003) as well as electron cryo-microscopy (Short et al., 2016; Xu et al., 2017) show that RAD51 assembly is mediated by protomer-protomer contacts that structurally mimic the RAD51-BRC repeat interaction. Because it interrupts these contacts *in vitro*, CAM833 is expected to suppress the function of RAD51 and prevent the formation of RAD51 foci in cells exposed to DNA damage.

We tested this prediction by monitoring RAD51 foci formation after the exposure of A549 non-small cell lung carcinoma (NSCLC) cells to ionising radiation (IR), using a robust cell-based assay based on high content microscopy with the Cellomics ArrayScan V^TI^, to objectively enumerate RAD51 foci (Jeyasekharan et al., 2013). IR-induced DNA breakage was monitored in the same experiment by enumerating foci containing γH2AX, a phosphorylated form of histone H2AX that is formed at DNA breaks (Rogakou et al., 1998).

Notably, CAM833 inhibited RAD51 foci formation 6 h after exposure to 3 Gy IR, in a concentration-dependent manner with an IC_50_ of 6 μM (Figure 3A and 3B, plotted as mean ± SEM, *n*=27). No RAD51 foci were detected at ∼50 µM CAM833, corresponding to a maximal level of inhibition. Furthermore, 50 µM CAM833 increased γH2AX foci formation 24 h after exposure by approximately 4-fold compared to control-treated cells (Figure 3C), suggestive of the persistence of unrepaired DNA damage. These findings are consistent with prior results in cells over-expressing BRC peptides (Chen et al., 1999b), and provide evidence that CAM833 engages its target in the cellular milieu to suppress RAD51 assembly and inhibit DNA repair by HDR.

**Figure 3.**
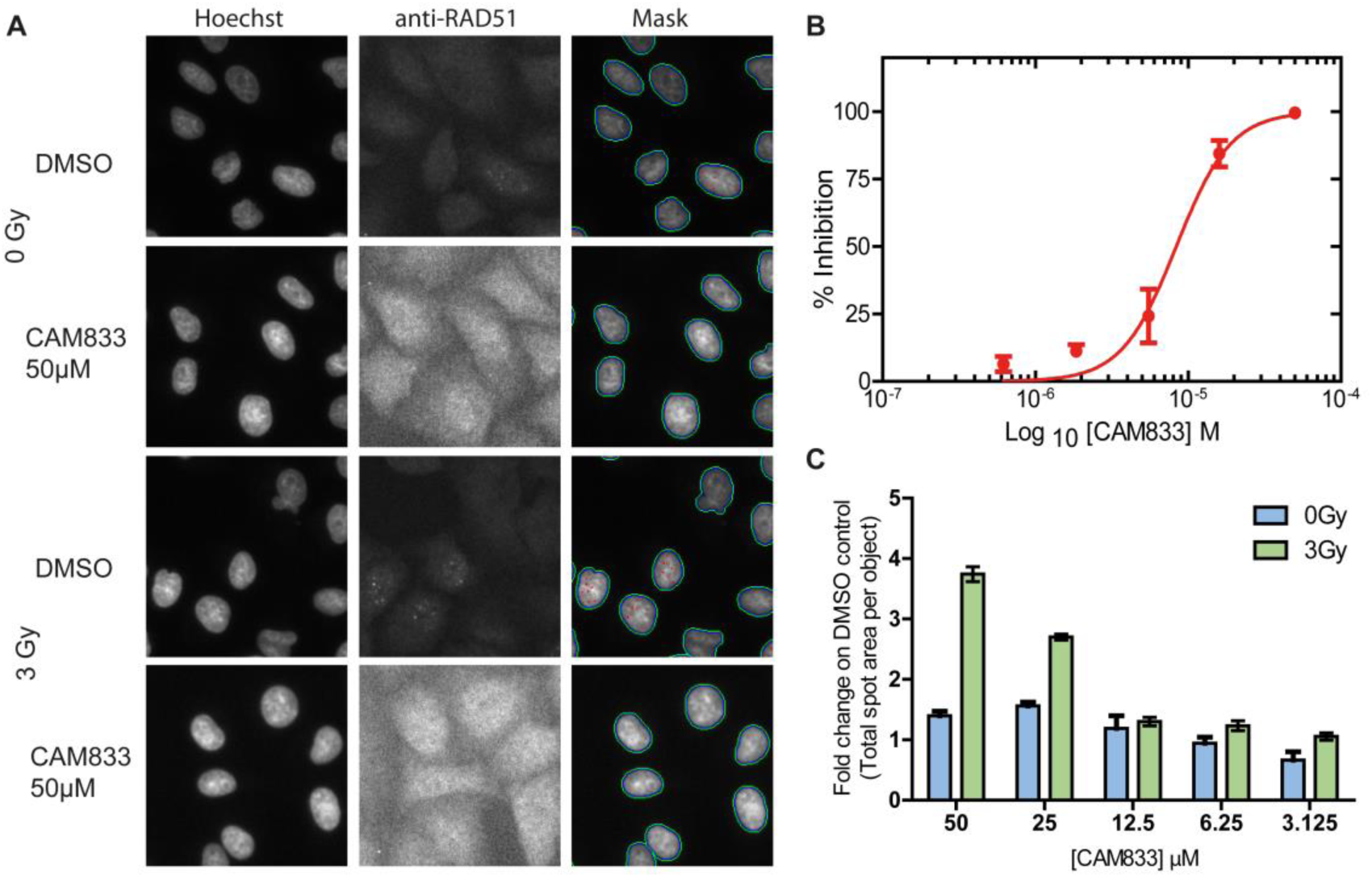
CAM833 causes a concentration-dependent decrease in RAD51 foci and subsequent increase in DNA damage. (A) Images from the Cellomics Arrayscan HCS microscope depicting A549 cells treated with CAM833 (50 µM) or DMSO controls with or without ionising radiation (3 Gy). Briefly, cells were co-stained with Hoechst-33342 to identify nuclei and anti-RAD51 antibody to detect RAD51 foci. The final column shows the Hoechst-stained cells with computationally identified nuclei outlined with green, and RAD51 foci with red, respectively. (B) An IC_50_ curve calculated from the images collected using the Cellomics HCS microscope as shown in 3A. A549 cells were treated with CAM833, exposed to 3 Gy ionizing radiation (IR) and fixed after 6 hours incubation. CAM833 inhibits the formation of IR induced RAD51 foci in A549 cells with an IC_50_ of 6 μM. Percent inhibition on the y-axis was plotted against CAM833 concentration (as log_10_M) on the x. Plots show mean ± SEM. (C) Cells treated by the same method were stained and counted for γ-H2AX foci 24 hrs after exposure. Each pair of bars corresponds to cells exposed to one of five different concentrations (lowest, 3.125 µM on the right, to highest, 50 µM, on the left) of CAM833 alone (0 Gy), or CAM833 plus 3 Gy IR (3 Gy). Bars depict the mean values of the fold change in γ-H2AX foci number over control cells treated with DMSO alone, ±SEM. CAM833 causes a concentration-dependent increase in unresolved DNA damage after 24 hours.

### CAM833 inhibits RAD51 molecular clustering after DNA damage

We have recently visualized the assembly of RAD51 molecules on DNA substrates at cellular sites of DNA damage using single-molecule localization microscopy (SMLM) by direct stochastic optical reconstruction (d-STORM) (Haas et al., 2018). Clusters of approximately 5-10 RAD51 molecules are first recruited to DNA damage sites 0.5-1 h after damage induction, which progressively extend into filaments >200 nm in length 3-5 h afterwards. SMLM shows that RAD51 clustering is suppressed by the over-expression of BRC repeat peptides, indicative of its dependence on protomer-protomer contacts that structurally mimic the RAD51-BRC repeat interaction inhibited *in vitro* by CAM833.

Therefore, to test the effect of CAM833 on RAD51 clustering we used SMLM on patient-derived EUFA423 cells (Figure 4A) bearing compound heterozygosity for the cancer-associated *BRCA2* truncating alleles 7691*ins*AT and 9000*ins*A (Haas et al., 2018; Howlett et al., 2002). We developed, as isogenic controls, EUFA423 cells complemented by the expression of full-length BRCA2 (EUFA423+BRCA2) (Hattori et al., 2011). We enumerated the number of RAD51 molecules detected by SMLM in clusters induced by the exposure of EUFA423 cells or EUFA423+ BRCA2 controls (Figure 4A) to 3 Gy IR, in the presence or absence of 25 µM CAM833, using a suite of bespoke image analysis algorithms that we have recently reported (Haas et al., 2018). As expected, the accumulation of RAD51 molecules in damage-induced clusters is significantly reduced in BRCA2-deficient EUFA423 cells compared to EUFA423+BRCA2 controls (Figure 4B) (Hattori et al., 2011). Notably, addition of 25 µM CAM833 significantly reduces RAD51 accumulation in damage-induced foci to a further extent in both cell types, providing additional evidence that CAM833 inhibits RAD51 protomer-protomer contacts during filament assembly.

**Figure 4.**
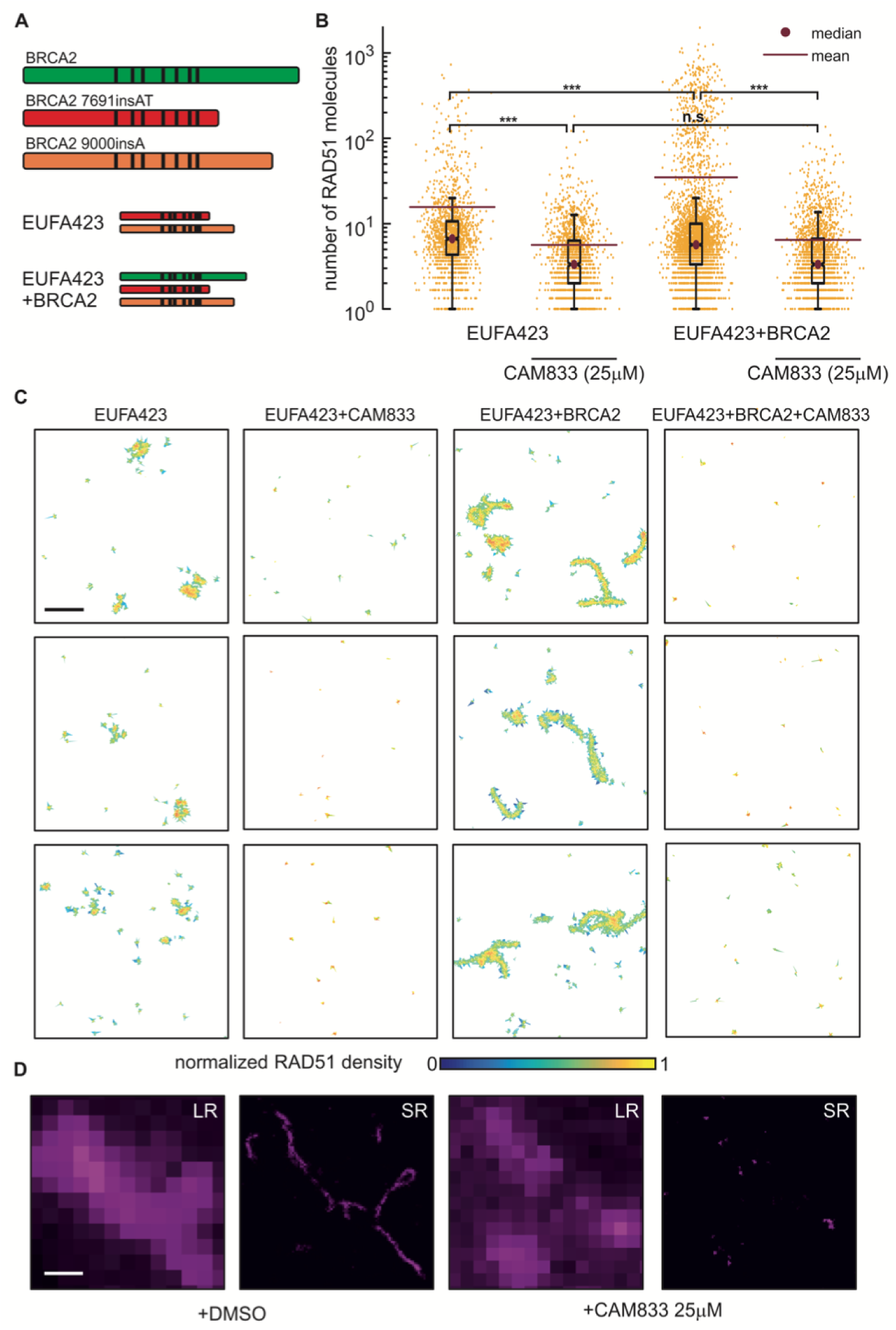
CAM833 inhibits RAD51 molecular clustering at DNA damage sites visualized by SMLM. (A) Diagrammatic representation of the bi-allelic truncating mutations (red and orange) affecting BRCA2 in the patient-derived cell line EUFA423, and their functional complementation by full-length BRCA2 (green) in EUFA423+BRCA2 cells. Black vertical lines depict the approximate positions of the BRC repeats. (B) Distribution of the number of RAD51 molecules contained within damage-induced clusters in EUFA423 or EUFA423+BRCA2 cells, without or with exposure to 25 µM CAM833, 3h after exposure to 3 Gy ionising radiation. Plots show the mean (purple line) ± standard error, as well as the median (purple dot) of the distributions. *** and n.s. indicates p-values lower than 10^−5^ and not significant differences, respectively. (C) Representative SMLM images of RAD51, represented as 2D Voronoi polygons. The colour of the polygons shows molecular densities normalized to the maximum value. Scale bar: 500 nm. (D) High magnification SMLM images of damage-induced RAD51 filaments in EUFA423+BRCA2 cells (DMSO-control left-hand panels), and their suppression by CAM833 (right-hand panels), under the same experimental conditions, at higher magnification. Scale bar, 200 nm. Images are shown either at low-resolution (LR) or super-resolved (SR).

The inhibitory effects of CAM833 are clearly observed by visualization of damage-induced RAD51 clusters as two-dimensional Voronoi polygons scaled to the maximum molecular density (Figure 4C). The compound effectively suppresses RAD51 clustering in both cell types, and in particular, prevents the formation of elongated filaments in control EUFA423+ BRCA2 cells. Example dSTORM pixel images (Figure 4D) further illustrate these effects, providing multiple lines of evidence for CAM833 target engagement and mechanism of action in cells.

### CAM833 potentiates radiation-induced cell cycle arrest and increases apoptosis over time

Genetic inactivation of RAD51 enhances cellular sensitivity to ionising radiation, accompanied by cell cycle arrest at the G2 checkpoint for DNA damage (Sonoda et al., 1998; Su et al., 2008). We hypothesized that similar effects would be triggered by the exposure of cells to CAM833. Indeed, when HCT116 colon carcinoma cells exposed to 20 μM CAM833 and 3 Gy IR were cell-cycle profiled by flow cytometry 4-72 h after exposure, we observed that treatment with CAM833 causes an increase in the percentage of cells with 4N DNA four hours after irradiation. Over time, there is a drop in the percentage of cells with 4N DNA in both treated and control groups. However, whereas in the control the percentage of cells in the apoptotic subG1 fraction remains below 5% throughout, in the compound-treated cells this rises progressively to peak at 15% at 48 hours (Figure 5A). Thus, these results suggest that treatment with CAM833 increases the progression of G2/M-arrested cells into apoptosis, as opposed to recovery.

**Figure 5.**
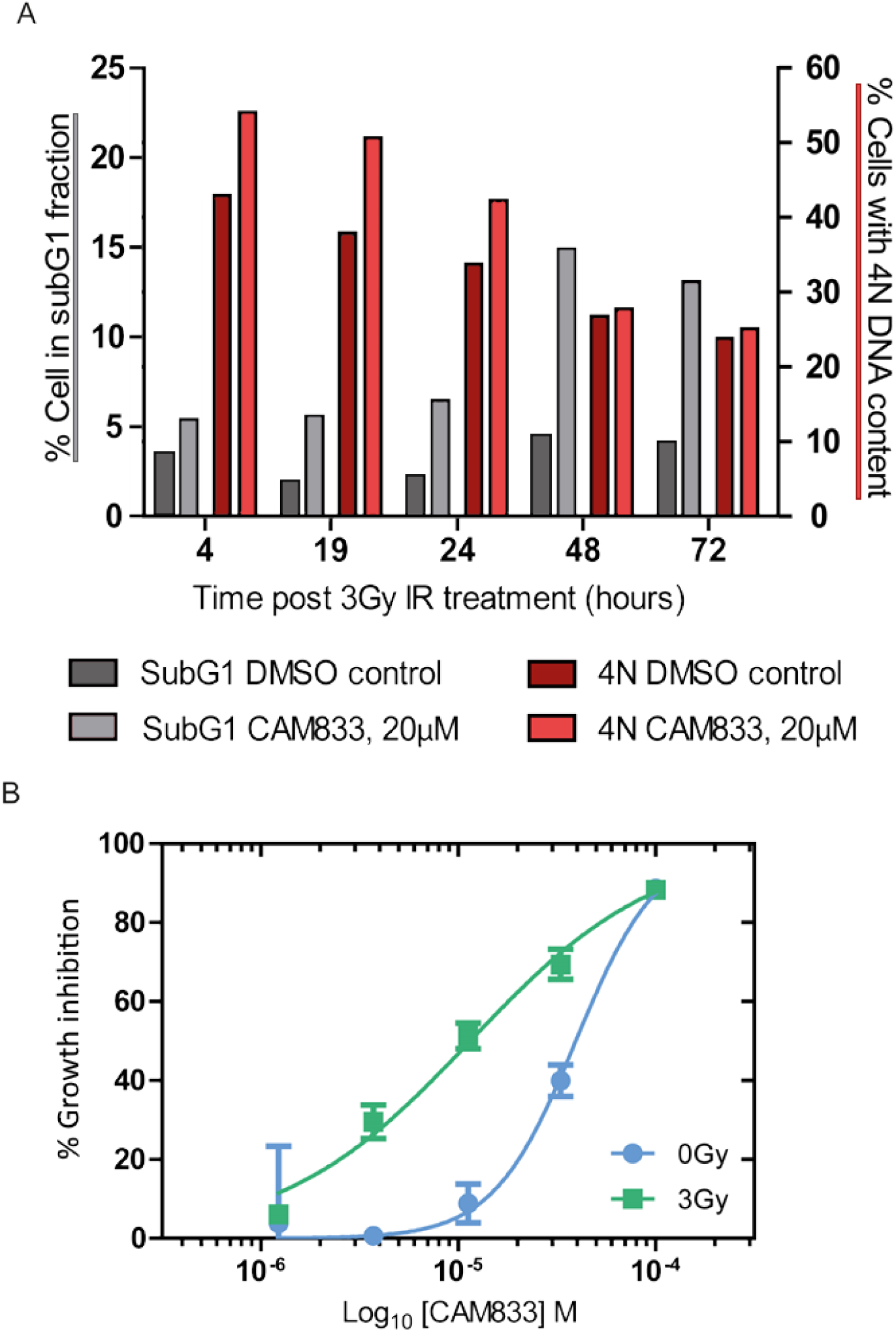
CAM833 potentiates radiation-induced cell cycle arrest with 4N DNA content and increased apoptosis over time. (A) Cell cycle analysis of HCT116 cells over a 72-hour time course after treatment with 20 μM CAM833 or DMSO control, combined with exposure to 3 Gy ionizing radiation. (B) plots the dose-response curves for growth inhibition of HCT-116 cells combining 0 (blue circles) or 3 Gy (green squares) of IR with different doses of CAM833 shown as log_10_M. Growth was measured after 96 hours using the sulforhodamine B cell proliferation assay. Each plotted value represents the mean percent growth inhibition ± SEM compared to control cells exposed to DMSO plus the indicated IR dose.

### CAM833 causes a dose-dependent growth inhibition which is enhanced when combined with ionising radiation

Consistent with these results, we find that CAM833 suppresses, in a concentration-dependent manner, the growth of multiple cancer-derived human cell lines (Supplementary Table 2). For instance, CAM833 alone inhibits the growth of HCT116 colon carcinoma cells with an average GI_50_ value of 38 µM (geometrical mean, *n*=18, SD 6.6 µM) after 96 h exposure. Moreover, our results suggest that CAM833 enhances cellular sensitivity to agents such as IR that induce DNA breakage normally repaired through RAD51-dependent HDR. Thus, when combined with 3 Gy IR, CAM833 suppresses the growth of HCT116 cells with a GI_50_ of 14 µM (geometrical mean, *n*= 18, SD, 6.2 µM), a concentration more than 2-fold lower than the GI_50_ for CAM833 alone (Figure 5B).

These findings prompted us to compare the effects of CAM833 with those of Carboplatin, a DNA cross-linking agent used in the clinic to sensitise cancers to therapeutic radiation (Clamon et al., 1999). We first exposed cells to a fixed 10 µM dose of either CAM833 or carboplatin, before treatment with 0-3 Gy IR, and compared cell growth using the sulforhodamine B cell proliferation assay 96 h afterwards (Figure 6A). Whereas carboplatin alone is more growth-suppressive than CAM833 alone, combination with increasing doses of IR potentiates the effects of CAM833 but not carboplatin (Figure 6A). The concentration-response curves (Figures 6B, 6C) showing the effect of combining 0-3 Gy IR with different concentrations of either carboplatin or CAM833 reflect a complex, dose-dependent response to the combined effects of CAM833 with IR, leading to changes in the observed IC_50_ (Figure 6D). IR at 1-2 Gy sharply potentiates the growth-inhibitory effects of 5×10^−5^ to 5×10^−4^M doses of CAM833. IR at 3 Gy has a smaller effect, across a wider dose range of CAM833. These differences could arise from biological factors such as variations in the amount or type of IR-induced DNA lesions, and/or the relative contribution of HDR to their repair. Collectively, these findings suggest the potential utility of CAM833 as a radio-sensitizer.

**Figure. 6.**
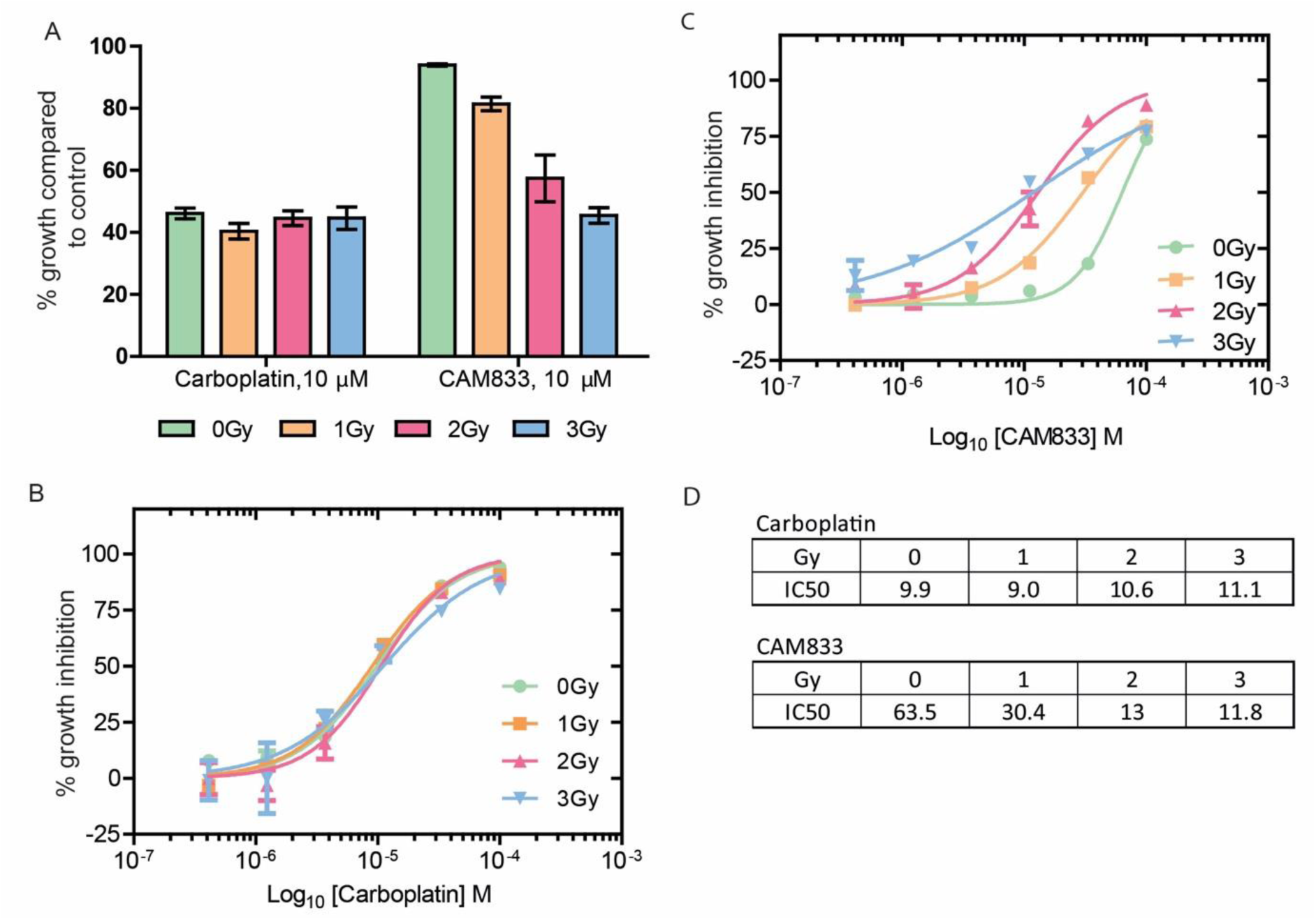
Low-dose ionising radiation potentiates the effects of CAM833 but not carboplatin. (A) Cell growth after exposure to increasing levels of ionizing radiation in the presence of a fixed dose (10 µM) of either Carboplatin or CAM833. Bars depict percent growth compared to control cells exposed to DMSO plus the indicated IR dose, represented as the mean ± SEM. Values < 100 indicate growth inhibition. (B) and (C) plot dose-response curves for growth inhibition combining 0 (green circles), 1 Gy (orange squares), 2 Gy (red triangles) or 3 Gy (blue triangles) of IR with different doses of carboplatin (B) or CAM833 (C) shown as log_10_M. In B-C, each plotted value represents the mean percent growth inhibition ± SEM compared to control cells exposed to DMSO plus the indicated IR dose. (D) shows the observed changes in IC_50_ (expressed in µM) for growth inhibition derived from the curves in (B) and (C). These data are representative of 3 independent experiments.

### CAM833 potentiates PARP1 inhibition in cells wildtype for BRCA2

Cells deficient in RAD51-mediated HDR through the inactivation of tumour suppressor genes like *BRCA1* or *BRCA2* exhibit hypersensitivity to poly-ADP ribose polymerase 1 (PARP1) inhibitors (Bryant et al., 2005; Farmer et al., 2005). We therefore tested whether CAM833 could potentiate the growth inhibitory effects of PARP1 inhibition by the inhibitor AZD2461 (Jaspers et al., 2013) in cells wildtype for BRCA2 (Figure 7). To this end, we determined dose-response curves for growth inhibition in cells exposed to different doses of AZD2461 combined with a fixed dose of either 10 µM (Figure 7A) or 20 µM (Figure 7B) of CAM833. While CAM833 alone had little effect (blue triangles), its combination with AZD2461 potentiated the growth-suppressive effects of PARP1 inhibition in a dose-dependent manner. Reciprocally, we also measured the dose-response curves for growth inhibition in cells exposed to different doses of CAM833 combined with a fixed dose of either 0.1 µM (Figure 7C) or 1 µM (Figure 7D) of AZD2461. These doses of AZD2461 have little effect when administered alone (blue diamonds), but again, their combination with CAM833 potentiates growth suppression by PARP1 inhibition in cells wild-type for BRCA2.

**Figure 7.**
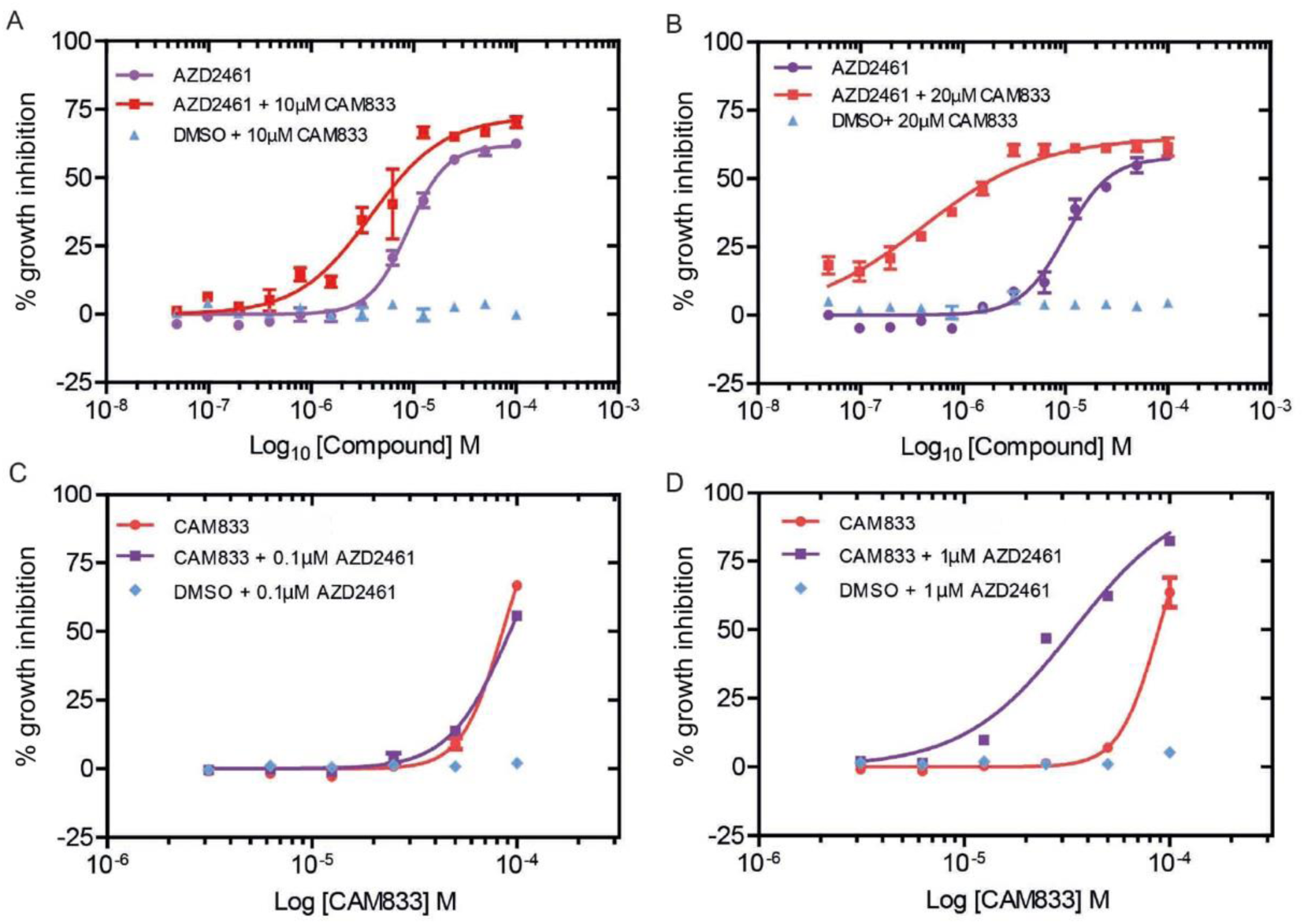
CAM833 potentiates the growth suppressive effect of PARP1 inhibition in BRCA2 wild-type cells. (A) and (B) show the dose-response curves for growth inhibition in HCT116 cells exposed to different doses of AZD2461 plotted as log_10_M combined with a fixed dose of either 10 µM (A) or 20 µM (B) of CAM833. Control experiments in which vehicle (DMSO) was added in place of AZD2461 are plotted in blue. Growth was measured 96 h after compound exposure using the SRB assay, and is depicted as the mean percent inhibition ± SEM compared to controls. (C) and (D) show reciprocal dose-response curves for growth inhibition after exposure to different doses of CAM833 plotted as log_10_M combined with a fixed dose of either 0.1 µM (C) or 1 µM (D) of AZD2461. Control experiments in which vehicle (DMSO) was added in place of CAM833 are plotted in blue. Measurements and plots are as in the previous panels.

## DISCUSSION

We report here the discovery of CAM833, a sub-micromolar chemical inhibitor of the regulatory protein-protein interaction between the RAD51 recombinase and the BRC repeat motifs of the tumour suppressor BRCA2. Using structure determination by X-ray crystallography, we show that CAM833 engages with two hydrophobic pockets on the surface of RAD51 that normally accommodate conserved hydrophobic side chains from the BRC repeats of BRCA2, thereby directly competing with the RAD51:BRCA2 interaction. These pockets also normally mediate RAD51 multimerization on DNA substrates during the process that leads to HDR, by accommodating corresponding hydrophobic residues from an adjacent RAD51 protomer to form the protomer-protomer interface. Consistent with these structural considerations, we show that CAM833 suppresses the assembly of RAD51 into damage-induced filaments visualized by single-molecule localization microscopy. Moreover, we present multiple lines of evidence suggesting that CAM833 potentiates growth inhibition, cell cycle arrest and cytotoxicity induced by DNA damage, consistent with its predicted ability to suppress DNA repair by HDR. Our findings have several important implications.

CAM833 is a well-characterized, selective chemical probe molecule which should prove valuable for further elucidating the biology of the RAD51-BRCA2 protein-protein interaction and the associated HDR pathways. Moreover, CAM833 is a chemically tractable starting point for the further, structure-guided development of optimized inhibitory compounds with the potential for development into a drug compound suitable for clinical studies. The development of this molecule through an innovative strategy of combining a fragment hit with a peptide lead compound reveals what is likely to be a generally-applicable strategy for the development of inhibitors of protein-protein interactions featuring a continuous peptide epitope (Scott et al., 2016).

Our work exemplifies a strategy to modulate the activity of RAD51 during HDR through two of its key regulatory protein-protein interactions. The first of these interactions is between RAD51 and the BRC repeats of BRCA2, which is essential to target RAD51 to cellular sites of DNA damage, and may also regulate RAD51 assembly on DNA substrates at these sites. The second interaction blocked by CAM833 is between RAD51 protomers, which occurs at the same structural motif engaged by the BRC repeats, and enables RAD51 assembly by multimerization. Our findings provide several lines of evidence that CAM833 acts in cells to engage RAD51 and block the protein-protein interactions that lead to its multimerization at sites of DNA damage. We find using SMLM by d-STORM that CAM833 suppresses the molecular clustering of RAD51 at damage sites, and prevents the extension of these clusters into extended RAD51 filaments, providing evidence for target engagement and the proposed mechanism of action. The mechanism of CAM833 action via the inhibition of RAD51-mediated HDR is further supported by our finding that the compound sensitizes cells with wildtype BRCA2 to the growth inhibitory effects of the PARP1 inhibitor, AZD2461. In the context of wildtype BRCA2, PARP1 inhibition alone is usually ineffective. While these results further support the cellular mechanism underlying CAM833 action, we are skeptical that systemic inhibition of RAD51 combined with the systemic effects of PARP1 inhibition has therapeutic potential owing to the likelihood of dose-limiting mechanism-related toxicity in normal tissues.

However, CAM833 also potentiates the cellular effects of ionizing radiation, a potent inducer of DNA breakage. When combined with IR, CAM833 sensitizes cells to IR-induced cell cycle arrest at the G2/M phase of the cell cycle, and enhances cell death by apoptosis. Collectively, these findings provide evidence supporting the further development of small-molecule inhibitors of the regulatory protein-protein interactions of RAD51 for cancer therapy through radiosensitisation.

## SIGNIFICANCE

Protein-protein interactions that mediate intracellular reactions leading to the repair of damaged DNA are an important target for anti-cancer drug discovery. Here, we report using structure-guided lead discovery the development of a potent orthosteric inhibitor, CAM833, of the protein-protein interaction between the BRCA2 tumour suppressor and the RAD51 recombinase, which is critical for the error-free repair of DNA breakage by homologous DNA recombination. The significance of our work is three-fold. First, it exemplifies a strategy for the development of inhibitors that target protein-protein interactions wherein a contiguous series of amino acids interact with a protein surface, by merging a peptidic inhibitor derived from those amino acids with chemical fragment hits identified by biophysical and crystallographic screening. Second, we demonstrate using single-molecule localization (“super-resolution”) microscopy that CAM833 inhibits RAD51 molecular clustering to prevent the assembly of extended RAD51 filaments at sites of DNA damage, validating target engagement, and demonstrating a unique mechanism of action. Finally, we show that CAM833 inhibits the cellular response to DNA damage, potentiating in BRCA2 wild-type cells the cytotoxic effects both of ionizing radiation or of PARP1 inhibitors, opening future avenues for anti-cancer drug development.

## Acknowledgements

We thank Dr Adrian Schreyer for developing a database for data management and Dr Tara Pukala and Prof Carol Robinson for taking part in the early stages of this project. We are grateful for Diamond Light Source for access to beamlines I02, I04 and I24 (proposals mx315 and mx7141) and ESRF for access beamline ID14-4. We thank the X-ray crystallographic and Biophysics facilities at the Department of Biochemistry for support and access. This work was funded by the Wellcome Trust Translational Award (080083/Z/06/Z) and Seeding Drug Discovery Award (91050/Z/10/Z) and we acknowledge the support of their Seeding Drug Discovery team, in particularly Dr Sarah Hardy and Prof Chas Bountra, for useful discussions. This work was also funded by Medical Research Council (MRC) Programme grants MC_UU_12022/1 and MC_UU_12022/8 to ARV, and a research grant from Astex Pharmaceuticals. We thank WuXi AppTec for Pharmacokinetic and cell-line sensitivity data and Cyprotex for ADMET data.

## Author contributions

Experimental investigations in Figs. 1, 2 were carried out by DES, TPS, AGC, GF, CV, TM, ARB, MEM, RS, DH, AH, ME, TP and JS; in Figs. 3, 5, 6 & 7, by NJF-N and LB; and in Fig. 4, by KH and AE. TLB, ARV, CA, MH, LP, GM, JS and TP conceptualised the project, supervised the experimental work, and analysed the results. DES, NJF-N, JS, MH, CA and ARV wrote the manuscript with input from all the authors. Each corresponding author supervised and is responsible for distinct aspects of this multi-disciplinary project. Chemistry was led by JS and CA; biochemistry and structural biology by MH; and microscopy, cell genetics and cell biology by ARV.

## Declarations

AV, LP and TB are inventors on a patent WO2004035621 - Use of crystal structure of human RAD51-BRCA2 repeat complex in screening for anti tumour agents.

## MATERIALS AND METHODS

### Chemical synthesis

See supplementary data for synthetic methods for all compounds.

### ITC

ITC was performed using a Microcal ITC-200 instrument at 25 °C. Experiments typically involved titrating a 10-fold excess of ligand in the injection syringe against the protein ([HumRadA2] = 60 µM or [ChimRAD51] = 20 µM) in either 200 mM Tris buffer at pH 7.5 and 100 mM NaCl (HumRadA2) or 20 mM potassium phosphate at pH 8.0 and 192 mM KCl (ChimRAD51). Titrations were typically performed with 5-10% DMSO and care was taken to ensure that the DMSO concentrations in the protein and ligand solutions were well matched. The raw ITC data were fitted using a single-site binding model in Microcal ITC LLC data analysis program in the Origin 7.0 package.

### FP assay

Fluorescence Polarisation (FP) competition experiments were performed as described in (Moschetti et al., 2016).In brief, binding of 10 nM Alexa Fluor 488-labelled BRC4 peptide to 50 nM ChimRAD51 protein (giving approximately 80-90 % saturation of binding) was competed with increasing concentration of inhibitor and the resulting competitive binding isotherms were measured and fitted using the expression described by (Wang, 1995).

### X-ray crystallography

Crystallisation and structure determination was done as described in (Moschetti et al., 2016). Ligands were soaked into HumRadA1 or HumRadA22F crystals in the presence of cryo-protectant typically overnight and crystals cryo-cooled in liquid N_2_. Diffraction data was collected at Diamond and ESRF synchrotrons and processed with XDS or autoproc (Kabsch, 2010; Vonrhein et al., 2011). Structures were solved by molecular replacement using corresponding apo structures and ligands fitted into the emerging density after brief refinement and complex structures refined to completion using phenix.refine or autoBuster (Adams et al., 2010). All crystallographic statistics are shown in Table S2 and coordinates and structure factors deposited in the Protein Data Bank under accession numbers 6TV4, 6TWR, 6TW4 and 6TW9.

### Cell culture

HCT116 colon carcinoma cells and A549 lung adenocarcinoma cells were obtained from ATCC and supplied mycoplasma free. HCT116 cells were maintained in McCoy’s 5A (1x) + Glutamax-I growth medium (Gibco, 36600-021) supplemented with fetal bovine serum (FBS, Gibco Life Technologies, 10270-106) at a final concentration of 10%. A549 cells were cultured in Dulbecco Modified Eagle medium (DMEM) (1x) +Glutamax-I (Gibco Life Technologies, 31966-021) with 10% FBS. All cells were grown at 37 °C/ 5 % CO_2_ in a humidified environment and all the assays were performed using these culturing conditions.

### Immunofluorescent visualisation of RAD51 foci/γH2AX foci in A549 cells using the Cellomics Arrayscan V^ti^ high content microscopy

A549 cells were seeded at 15000 cells/well in 100 μl (1.5×10^5^ cells/ml) in Nunc 96-well plates (cat# 167008) and grown overnight prior to the drug treatment. Compounds were added to cells such that the final DMSO concentration did not exceed 1% v/v. Following compound addition, cells were exposed with specified levels of ionising radiation using the Xstrahl RS225 X-ray generator. After incubation with the compound for 6 hours, the medium was removed by aspiration and the cells washed twice in 1xPBS. Cells were fixed using fixative solution (4% formaldehyde diluted in PBS) pre-warmed to 37°C for 10 min at room temperature. Cells were then washed twice in 100 μl 1x PBS at room temperature. Cells were then incubated in 100μl permeabilisation buffer for 5 minutes at room temperature after which they were incubated with 100μl of blocking buffer (2% BSA (w/v), 0.2% Tween v/v, 0.1% TritonX-100 (v/v) in PBS) for 90 minutes at room temperature. Cells were subsequently incubated with 50 µl of mouse polyclonal anti-RAD51 Antibody (Abnova, cat # H00005888-B01P) diluted 1:200 in blocking solution for 2 h at room temperature. Cells were washed in 100 µl wash buffer at room temperature (0.2% Tween (v/v), 0.1% Triton X-100 (v/v) in 1x PBS) then incubated in 50 µl Alexa Fluor 488 labelled anti-mouse secondary antibody (1:500) and Hoechst 33342 (10 mg/ml stock) counterstain at 1:1000 in blocking solution for 60 minutes at room temperature. Finally, cells were washed twice in wash buffer and then twice in PBS and then stored in 100 µl in PBS with a light protective seal at 4 °C until read on the Cellomics Arrayscan V^ti^ using a spot detector protocol. The number of cells analysed was 800 and the parameter used for analysis was Total Spot Area.

For detection of γH2AX foci in A549 cells, 10,000 cells/well (1×10^5^ cells/ml) were seeded in 100 μl and left to grow overnight before treatment with compound. Cells were subsequently exposed to compounds and either 3 Gy ionising radiation or mock treated (left on the bench at room temperature). Staining protocol was identical as for RAD51 foci but anti-phospho γH2AX primary mouse monocolonal antibody was used (Milipore, cat#05-636) at 1:2000 dilution.

### SRB growth inhibition assay

Adherent cell lines (HCT116 and A549 cells) were seeded into flat-bottomed tissue culture 96-well plates in a volume of 150 µL of growth medium. HCT116 cells were seeded at 750 cells per well and A549 cells were seeded at 1000 cells per well. After 24 hours, compounds dissolved in DMSO were diluted in growth medium and were added to cells such that the final DMSO concentration was 1% (v/v) and the final volume in the well was 200 µL. Cells were then incubated in the presence of compound for 96 hours before fixation.

Medium was removed from cells and 100 µL of cold 1% (v/v) trichloroacetic acid was added for 30 minutes at 4 degrees. The plates were washed three times in tap water and left to dry at room temperature. The fixed cells were stained in a 0.057% sulphorodamine B/1% acetic acid solution (w/v) and incubated at room temperature with agitation for 30 minutes after which the dye was removed and the plates washed in 1% (v/v) acetic acid and left to dry. The dye was then solubilised in 10 mM Tris solution (pH8) and incubated for 30 minutes under agitation. The plates were then read on a PHERAstar plus plate reader (BMG Labtech) using the fluorescence intensity module (540-590 nm). Growth inhibition was calculated relative to DMSO controls and GI_50_ values were calculated using Graphpad Prism.

For the PARP inhibitor experiments, the SRB method was used as described above to measure growth inhibition with the exception that cells were seeded into 150 µl medium and then a combination of either 25 µl of CAM833, AZD2461 or DMSO was added to give a total volume of 200ul in the well.

### Flow cytometry

Propidium iodide staining solution (PI solution) was used at the following final concentrations: 200 ug/ml RNAase A (Sigma Aldrich, cat# 10109169001), 0.1% Triton-X 100 and 20 ug/ml of propidium iodide solution diluted in 1x PBS. HCT116 cells were grown in 6-well plates in a total volume of 2 ml and treated with either test compound or DMSO control for the designated time. After treatment, medium was collected from the cells which were then washed in 1x PBS then removed from the plastic by the addition of in 500 μl Trypsin/EDTA until cells were monodispersed. The trypsin was neutralised by the removed media and the cell suspension was spun at 1000 rpm for 5 minutes. Cells were then washed a further time in ice cold 1xPBS and spun at 1000 rpm for 5 minutes. Cells were then fixed in 4.5ml 70% ice cold ethanol and 0.5 ml ice cold 1xPBS. Cells were left in fixing solution overnight at 4 ^°^C until processing. Cells were spun at 1000 rpm for 5 mins and then washed in 1xPBS, re-suspended in 0.5-1 ml of the PI solution at incubated in the dark for 2 hours at room temperature. Cells were then counted and analysed using a Becton Dickinson LSR II cytometer and FCS Express software.

### Super-resolution microscopy

Single Molecule Localization Microscopy (SMLM) was achieved by direct Stochastic Optical Reconstruction Microscopy (d-STORM) as described (Haas et al., 2018). Briefly, samples were prepared for one colour 2D d-STORM utilizing a buffer containing 100 mM MEA-HCL (Sigma, M6500), 10% glucose (Sigma), 0.5 mg/ml glucose oxidase (Sigma, G2133) and 40 µg/ml catalase (Sigma, C100) in water at pH 7.5. Samples were imaged by direct STORM at room temperature in sealed 8-well ibidi µ-slides utilizing an inverted N-STROM microscope (Nikon Ti, Japan) equipped with an Apochromat 100x/1.49 NA oil immersion objective. Samples were let to equilibrate for at least 30 minutes before imaging to minimize thermal drift. Images were then acquired with highly inclined illumination and focus was maintained by hardware autofocusing (Nikon Perfect Focus System). AlexaFluor647 was first pumped in its dark state using the 640 nm laser line at maximum power (∼150 mW) and then imaged continuously with a power density of ∼3 kW/cm^2^. Data were acquired in ‘streaming mode’ with a field-of-view (FOV) of 256×256 pixels (160 nm pixel size), at 65 frames per second for 25,000 frames with an EMCCD camera (iXon Ultra DU897, Andor). The sparsity of single molecules per frame was controlled using ∼0.6 mW of the 405 nm laser. Images of AlexaFluor647 were acquired with a Quad Band Set for TIRF applications (Chroma, TRF89901, ET – 405/488/561/640 nm) and the ET645/75 m emission filter (Chroma).

### Cluster data analysis

Single molecule data was analysed using the Grafeo program available at https://github.com/inatamara/Grafeo-dSTORM-analysis-, as described in (Haas et al., 2018). Briefly, all localizations with fewer than 1,000 detected photons or localization precision lower than 20 nm were discarded. Next, the data was filtered using 2D Voronoi diagrams, setting the minimum density (an inverse of Voronoi polygon VP size) to 5*10^−5^ nm^-2^. Finally, small isolated detections were supressed by thresholding univariate distance distribution function – a detection was rejected if it had less than 20 neighbours at the distance ≤100 nm. Next, two-dimensional Delaunay triangulation (DT) was computed. Localizations were assigned to discreet clusters, connected components, by removing all DT edges larger than 20 nm. All segmented connected components having less than 3 localizations were discarded. The number of RAD51 molecules inside a cluster was estimated by dividing the number of localization within a cluster by the expected number of localization obtained from isolated secondary antibodies used to label RAD51.

### Statistical tests

Simultaneous comparisons of the median values of multiple groups were performed using the Kruskal-Wallis test at the significance level alpha of 0.05 and familywise error rate was corrected by adjusting p-values using the Tukey-Kramer method.

