## Supplemental data and methods for "A small-molecule inhibitor of the BRCA2-RAD51 interaction modulates RAD51 assembly and potentiates DNA damage-induced cell death"

**Figure S1.** ITC data for the binding of **3** (left, $K_d$ 1.36 mM) and **4** (right, $K_d$ 3 µM) to HumRadA2

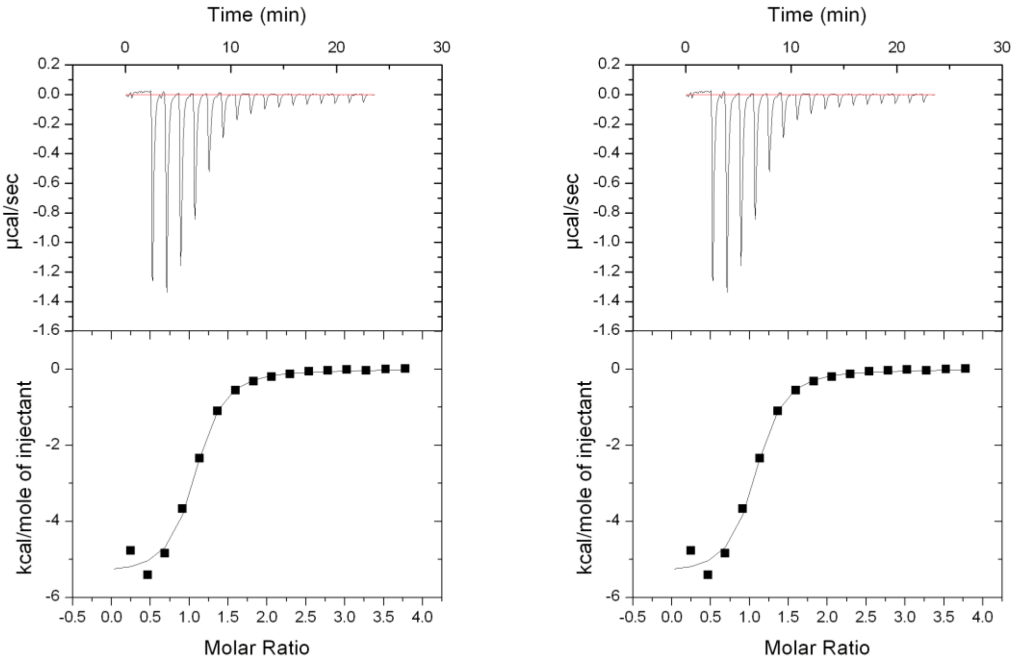

**Figure S2.** ITC data for the binding of **5** to HumRadA2 ($K_d$ 220 nM)

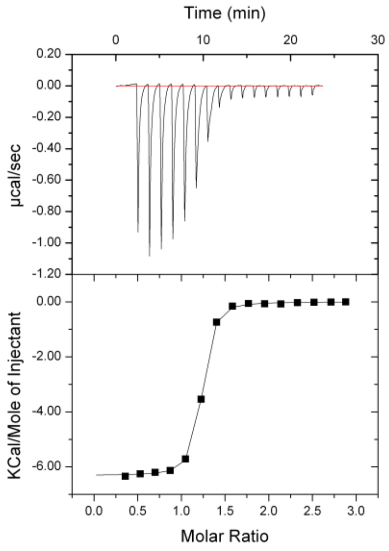

Figure S3. Structure of **6** bound to two forms of humanised RadA.

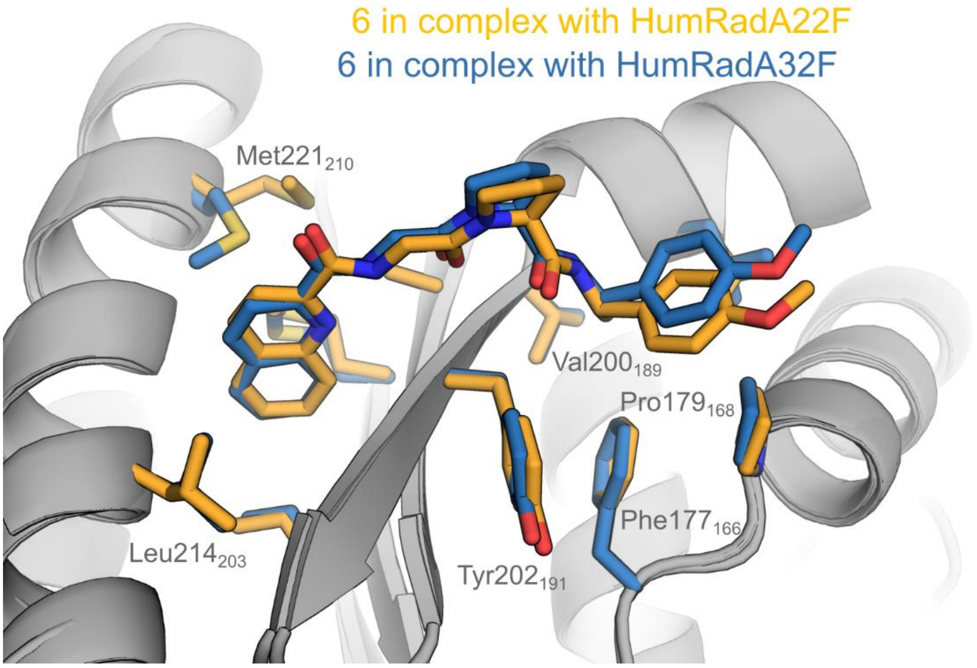

HumRadA22F (orange carbons, PDB:6TW4) and HumRadA32F (blue carbons, PDB: 6XTW) in complex with **6** with the side chains of residues around the Phe and Ala pockets shown as sticks and in the same colour as the ligand bound to that protein.

Figure S4. Inhibition of RAD51 oligomerisation by CAM833

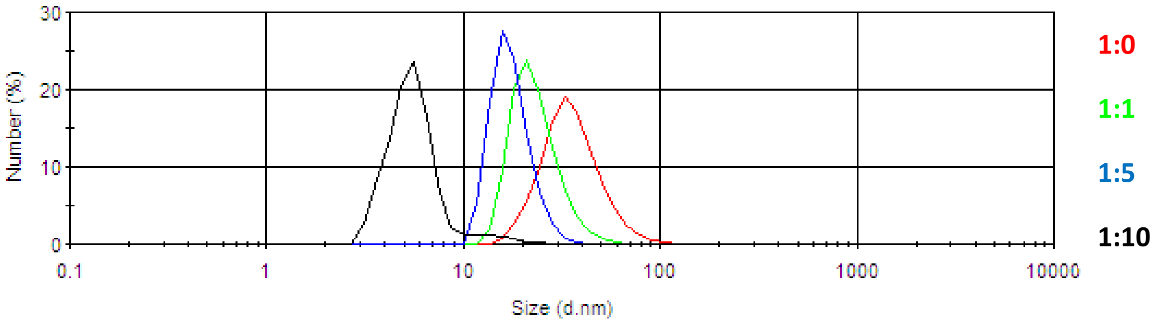

Particle size distribution (in nm) of full-length RAD51 protein in the absence (red line) and presence of increasing concentrations of CAM833. Green line indicates 1:1 stoichiometry of RAD51:CAM833, blue line is for sample with 1:5 stoichiometry and the black line for 1:10 stoichiometry of the components.

Figure S5. Structure of CAM833 bound to HumRadA22F.

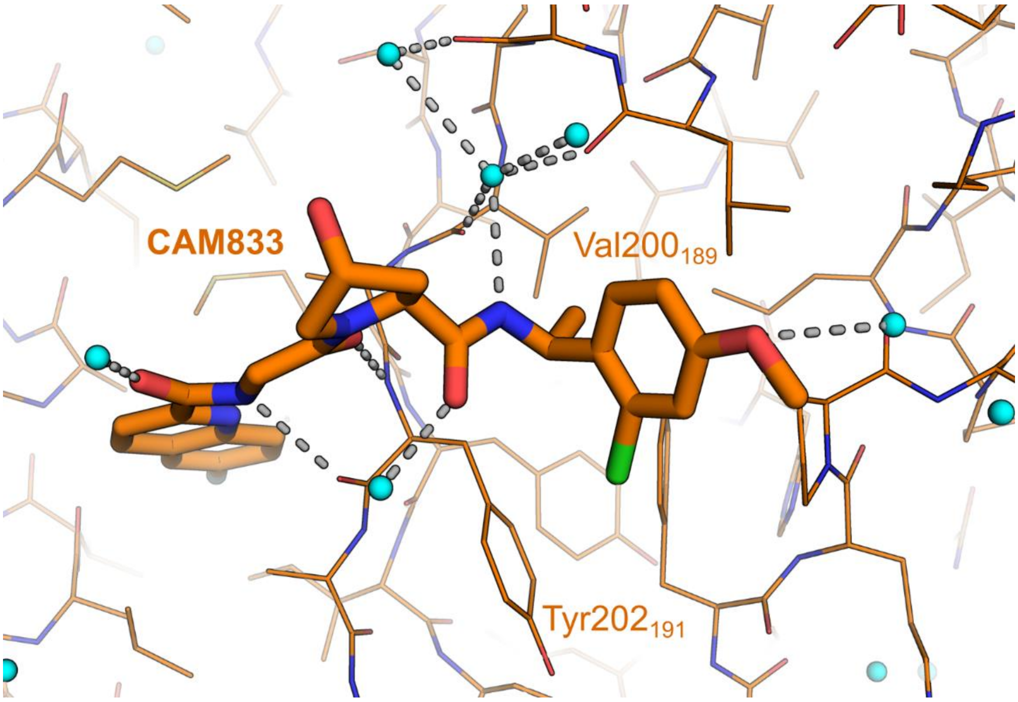

CAM833, in thicker sticks, is bound to HumRadA22F with hydrogen bonds between the inhibitor and the protein (thinner lines) or waters (small cyan spheres) shown as grey dotted lines.

Table S1. Calculated and measured ADMET and developability properties for CAM833.

| | |
|---|---|
| MW | 529 |
| PSA | 120 |
| clogP[a] | 2.73 |
| $K_D$ FP (ChimRAD51) | 355 nM,[b] (n=8) |
| Aqueous solubility [c] | 7.75 µg/ml (n=2, s.d. 0.549) |
| Microsomal $CL_{int}$ [d] | mouse 32.2 ± 2.99 µL/min/mg protein<br>rat 28.9 ± 4.16 µL/min/mg protein<br>human 18 ± 4.52 µL/min/mg protein |
| CYP450 $IC_{50}$ (1A2, 2C9, 2C19, 2D6, 3A4 midazolam and testosterone sites) | all > 25 µM |
| hERG electrophysiology @ 10 uM[e] | 12% |
| Human plasma protein binding [f] | 99.2% bound |
| Caco-2 permeability<br>AtoB $P_{app}$ ($10^{-6}$ cm $s^{-1}$) / Efflux Ratio | 0.23 / 195 |
| Selectivity Cerep ExpresSPanel | no inhibition of binding >50% at 10 µM |
| Mouse iv pharmacokinetics at 1 mg/kg[g] | t½ = 1.44 h (n=2)<br>CL 44.6 ml/min/kg (n=3)<br>$Vd_{ss}$ 2.51 L/kg (n=2) |
| Mouse oral screen at 50 mg/kg [h] | Cmax 8323 ng/ml (n=3)<br>F = 44% |

a. Calculated using Chemdraw 16

b. $pK_d$ = 6.45 ± 0.16, n=8 determined by FP

c. Thermodynamic solubility from solid, determined overnight in pH 7.4 buffer.

d. Intrinsic clearance calculated from 5 timepoints over a 45 minute experiment ± standard error

e. Inhibition of hERG tail-currents measured by whole-cell voltage-clamping in mammalian cells.

f. Determined by equilibrium dialysis

g. Fasted male CD-1 mice. 0.5 mg/mL in 20%HP-β-CD in water, clear solution. n=3 or 2

h. Fasted male CD-1 mice, fasted. 5 mg/mL in 70% PEG400 / 30% water, clear solution. n=3 or 2.

**Table S2.** CAM833 growth inhibition data for a range of cancer-derived human cell lines.

| Cell Line | Absolute $EC_{50}$ (µM) |
|---|---|
| HUVEC | 133.0 |
| Mia-pa-ca2 | 68.6 |
| NCI-H209 | 69.3 |
| OVCAR3 | 53.6 |
| PC3 | 91.2 |
| SK-MEL-24 | 143.5 |
| U20S | 40.5 |
| BT549 | 73.8 |
| HCT116 | 63.5 |
| A375 | 72.0 |
| BT20 | 61.9 |
| HT1367 | 104.0 |
| MDA-MB-453 | 48.0 |
| RT112/84 | 70.6 |
| T.Tn | 55.8 |
| U118MG | 90.3 |
| SCaBER | 47.0 |
| SK-OV-3 | 104.8 |
| T24/83 | 96.7 |
| U87 | 108.8 |
| A549 | 39.0 |
| Calu3 | 83.3 |
| HepG2 | 68.1 |
| HFL-1 | 98.8 |
| HL-60 | 87.7 |
| K562 | 137.1 |
| LN229 | 117.3 |
| Raji | 95.6 |
| RKO | 63.5 |
| 5637.00 | 84.5 |
| A2780 | 46.5 |
| Caki1 | 73.0 |
| PANC1 | 96.0 |
| SW756 | 124.3 |

Table S3. Crystallographic data collection and refinement statistics

| Ligand | **3** | **4** | **6** | **6** | **7 / CAM833** |
|---|---|---|---|---|---|
| Protein form | HumRadA1 | HumRadA1 | HumRadA22F | HumRadA33F | HumRadA22F |
| PDB code: | 6TV3 | 6TWR | 6TW4 | 6XTW | 6TW9 |
| **Data Collection and Processing:** | | | | | |
| Synchrotron beamline | ESRF ID14-4 | DLS I04 | DLS I24 | DLS I03 | DLS I02 |
| Wavelength (Å) | 0.9795 | 0.9702 | 0.9686 | 0.9300 | 0.9795 |
| Resolution range (Å)<br>(High resolution bin) | 21.03 - 1.50<br>(1.59 - 1.50) | 31.63 - 1.35<br>(1.43 - 1.35) | 40.39 - 1.73<br>(1.77 - 1.73) | 100.82-2.31<br>(2.43-2.31) | 61.21 - 1.52<br>(1.522 - 1.517) |
| Space group | P 1 21 1 | P 21 21 21 | P 21 21 21 | P31 2 1 | P 21 21 21 |
| Unit cell (a b c) (Å) | 37.67 79.24 39.43 | 40.53 61.95 87.72 | 40.39 60.15 88.20 | 89.83 89.83 100.82 | 40.38 61.21 87.68 |
| Unit cell ($\alpha$ $\beta$ $\gamma$) (°) | 90.00 118.18 90.00 | 90.00 90.00 90.00 | 90.00 90.00 90.00 | 90.00 90.00 120.00 | 90.00 90.00 90.00 |
| Total number of reflections | 104241 (31919) | 325400 (44541) | 182953 (12814) | 107445 (16048) | 149322 (1612) |
| Number of unique reflections | 31919 (5063) | 48841 (7575) | 23054 (1639) | 21180 (3039) | 32571 (351) |
| Multiplicity | 3.7 (3.6) | 6.7 (5.9) | 7.9 (7.8) | 5.1 (5.3) | 4.6 (4.6) |
| Completeness (%) | 97.4 (96.4) | 99.5 (96.9) | 99.6 (98.6) | 99.9 (99.9) | 94.6 (97.5) |
| Mean I/sigma(I) | 15.9 (2.9) | 13.8 (2.2) | 15.2 (0.6) | 14.4 (2.8) | 12.5 (2.2) |
| $R_{merge}$ | 0.049 (0.45) | 0.080 (0.64) | 0.087 (1.25) | 0.067 (0.621) | 0.087 (0.67) |
| $R_{pim}$ | - (-) | - (-) | 0.035 (0.50) | 0.036 (0.330) | 0.043 (0.34) |
| CC-half | 0.999 (0.87) | 0.998 (0.77) | - (-) | - (-) | - (-) |
| **Refinement:** | | | | | |
| R / $R_{free}$ | 0.183 / 0.217 | 0.187 / 0.210 | 0.178 / 0.210 | 0.184 (0.229) | 0.156 / 0.172 |
| Number of atoms | 1985 | 2334 | 1955 | 3664 | 3853 |
| No. of ligand atoms | 40 | 75 | 35 | 35 | 44 |
| No. of waters | 221 | 358 | 143 | 118 | 258 |
| No. of protein residues | 219 | 227 | 226 | 227 | 224 |
| Average/Wilson<br>B-factor (Å$^2$) | 24.5 / 18.5 | 18.2 / 19.3 | 27.8 / 23.2 | 48.0 | 17.0 / 15.2 |
| B-factor for ligands (Å$^2$) | 34.8 | 26.4 | 28.6 | 52.0 | 19.1 |
| B-factor for solvent (Å$^2$) | 39.3 | 29 | 39 | 46.9 | 30.6 |
| RMS (bonds) (Å) | 0.01 | 0.005 | 0.01 | 0.007 | 0.011 |
| RMS (bond angles) | 1.12 | 0.81 | 1.09 | 0.862 | 1.16 |

### Supplementary Experimental procedures – chemical synthesis

#### Solvents and Reagents

Unless otherwise stated starting materials and reagents were purchased from regular suppliers. Dry solvents were purchased and used as provided.

#### Chromatography

Thin layer chromatography (TLC) was performed on glass plates coated with Merck 60 F254 silica and visualization was achieved by UV light or by staining potassium permanganate. Flash column chromatography was using a Biotage Isolera One and Biotage Isolera Four systems with UV detection at 254 nm and 280 nm and commercially available cartridges.

#### Nuclear Magnetic Resonance Spectroscopy

$^1$H NMR spectra were recorded on a Bruker Avance 400 (400 MHz), or Bruker Avance Cryo 500 (500 MHz). Chemical shifts are quoted in ppm and are referenced to the residual non-deuterated solvent peak, and are reported (based on appearance rather than interpretation) as follows: chemical shift δ /ppm (multiplicity, coupling constant J/Hz, number of protons) [br, broad; s, singlet; d, doublet; t, triplet; q, quartet; qui, quintet; sept, septet; m, multiplet]. All J values are given in Hz. Fractional integrations are reported where conformational restriction of peptidic compounds results in separate signals on the nmr timescale.

#### LCMS

High-resolution mass measurements were performed on a Waters LCT Premier mass spectrometer or a Kratos Concept mass spectrometer. Low-resolution measurements were recorded on a Waters / ZQ LCMS and on a Waters Acquity UPLC HClass LCMS. All final compounds used for screening and cell experiments were at least 95% pure as determined by LCMS unless otherwise stated.

#### Synthetic methods

Tetrapeptide (**1**) was prepared as described previously (Scott et al., 2016). Fragments 2-naphthol (**2**) and 3-amino-2-naphthoic acid (**3**) are commercially available and used as supplied. Tetrapeptide **(4)** was prepared using standard solid phase chemistry with Fmoc protection by the Protein and Nucleic Acid Service at the Department of Biochemistry (University of Cambridge).

#### General procedure A

EDAC (1.5 equiv.) was added to a stirred solution of acid (1 equiv.), amine (2 equiv.), *N*-methylmorpholine (3 equiv.) and DMAP (1 equiv.) in DCM (0.05 M) and stirred until the reaction was judged complete. The solvent was removed *in vacuo* and DCM and $H_2O$ were added to the residue, the organic layer was separated, washed with $H_2O$, brine, dried with $MgSO_4$, filtered and the solvent removed *in vacuo*. The residue was purified by flash chromatography (FC) (2-20% MeOH: DCM) to give the product.

#### General procedure B

To a solution of acid (1 equiv.), amine (1-1.1 equiv.) and DIPEA (2-5 equiv.) in solvent was added PyBOP (1.1-1.4 equiv.) and the reaction was stirred until complete and then concentrated *in vacuo*. Unless stated otherwise, the crude was diluted with ethyl acetate and the organics were washed three times with water, and then brine and dried before purifying typically by FC as described to give the product.

(*S*)-1-((2-Naphthoyl)-L-alanyl)-*N*-((*S*)-1-phenylethyl)pyrrolidine-2-carboxamide (5)

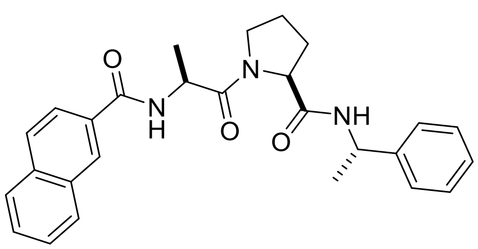

**Methyl (2-naphthoyl)-L-alanyl-L-prolinate**

A mixture of methyl L-alanyl-L-prolinate hydrochloride (200 mg, 0.84 mmol), 2-naphthoic acid (218 mg, 1.27 mmol), EDAC (243 mg, 1.27 mmol), *N*-methylmorpholine (204 µL, 1.86 mmol) and DMAP (103 mg, 0.84 mmol) in DCM (10 mL) was stirred until judged complete and purified by FC (1-10% MeOH: DCM) to give the product as a white foam (240 mg, 81%).

$^1$H NMR (500 MHz, Methanol-$d_4$) δ 8.43 (d, *J* = 1.6 Hz, 1H), 7.98 (dd, *J* = 7.8, 1.6 Hz, 1H), 7.96 – 7.88 (m, 3H), 7.62 – 7.54 (m, 2H), 4.52 (dd, *J* = 8.6, 4.7 Hz, 1H), 3.94 (dt, *J* = 9.9, 7.1 Hz, 1H), 3.75 (dt, *J* = 10.1, 6.6 Hz, 1H), 3.72 (s, 3H), 2.36 – 2.24 (m, 1H), 2.09 (p, *J* = 6.7 Hz, 2H), 1.99 (dtd, *J* = 12.5, 6.7, 4.8 Hz, 1H), 1.51 (d, *J* = 7.1 Hz, 3H). Proline α-CH is under the solvent peak (4.83-4.88 ppm)

LCMS m/z 355.3 $(M+H)^+$

**(2-Naphthoyl)-L-alanyl-L-proline**

To a solution of methyl (2-naphthoyl)-L-alanyl-L-prolinate (230 mg, 0.65 mmol) in THF:$H_2O$ (1:1, 20 mL) was added dropwise at 0 °C a solution of NaOH (104 mg, 2.6 mmol) in $H_2O$ (1 mL). The reaction mixture was stirred for 10 mins at 0 °C, allowed to warm to rt and stirred for 2 hours. The mixture was concentrated *in vacuo*, acidified on ice to pH 2 and the resulting white suspension was extracted with DCM (4 x 100 mL). The organic extracts were combined, dried with $MgSO_4$, filtered and the solvent removed in vacuo to give the acid as a white foam (212 mg, 96%) which was used without further purification.

$^1$H NMR (400 MHz, Chloroform-*d*) δ 8.30 – 8.24 (m, 0.85H), 8.23 (d, *J* = 1.3 Hz, 0.15H), 7.88 – 7.72 (m, 4H), 7.61 (d, *J* = 7.6 Hz, 0.15H), 7.53 – 7.43 (m, 2H), 7.38 (d, *J* = 7.6 Hz, 0.85H), 4.98 (p, *J* = 7.0 Hz, 0.85H), 4.85 (p, *J* = 6.8 Hz, 0.15H), 4.56 (dd, *J* = 7.0, 5.7 Hz, 0.85H), 4.49 (dd, *J* = 7.5, 3.2 Hz, 0.15H), 3.84 – 3.46 (m, 3H), 2.24 (td, *J* = 7.1, 3.1 Hz, 0.15H), 2.17 – 2.11 (m, 1H), 2.07 – 1.93 (m, 1.7H), 1.90 – 1.81 (m, 0.15H), 1.81 – 1.75 (m, 1H), 1.46 (d, *J* = 6.9 Hz, 2.55H), 1.42 (d, *J* = 6.7 Hz, 0.45H).

LCMS m/z 341.2 $(M+H)^+$

**(*S*)-1-((2-Naphthoyl)-L-alanyl)-*N*-((*S*)-1-phenylethyl)pyrrolidine-2-carboxamide (5)**

(2-Naphthoyl)-L-alanyl-L-proline (25 mg, 0.07 mmol) was dissolved in DCM (2 mL) and cooled to 4 ˚C under $N_2$ gas. To the stirred solution was added EDAC (21 mg, 0.11 mmol), (S)-α-methylbenzylamine (19 µL, 0.15 mmol) and DMAP (10 mg, 0.07 mmol). The reaction was stirred for 3 hours, and the solvent was removed *in vacuo.* The residue was dissolved in ethyl acetate (100 ml), washed with water (2 x 100 ml), washed with brine and then dried ($MgSO_4$) and the solvent was removed *in vacuo*. The crude product was purified by FC (1-10% MeOH: DCM) to give the product as a clear oil (18 mg, 54%).

$^1$H NMR (400 MHz, Chloroform-*d*) δ 8.27 (s, 1H), 7.90 – 7.82 (m, 1H), 7.82 – 7.77 (m, 3H), 7.48 (ddt, *J* = 8.0, 6.9, 5.3 Hz, 2H), 7.33 – 7.13 (m, 6H), 7.04 (d, *J* = 8.1 Hz, 1H), 5.06 – 4.87 (m, 2H), 4.54 (dd, *J* = 8.2, 2.8 Hz, 1H), 3.71 (td, *J* = 9.1, 8.6, 7.2 Hz, 1H), 3.59 (ddd, *J* = 9.8, 8.0, 4.1 Hz, 1H), 2.37 – 2.24 (m, 1H), 2.22 – 2.07 (m, 1H), 2.05 – 1.89 (m, 1H), 1.91 – 1.74 (m, 1H), 1.46 (d, *J* = 6.8 Hz, 3H), 1.39 (d, *J* = 6.9 Hz, 3H).

LCMS m/z 442.1 $(M-H)^-$

*N*-(2-((*S*)-2-(((*S*)-1-(4-methoxyphenyl)ethyl)carbamoyl)pyrrolidin-1-yl)-2-oxoethyl)quinoline-2-carboxamide (6)

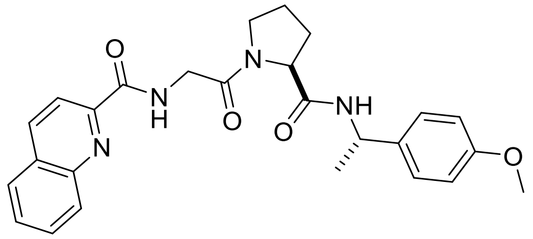

**(Quinoline-2-carbonyl)glycyl-L-proline**

To a mixture of (quinoline-2-carbonyl)glycine (1.9 g, 8.26 mmol) and L-proline-t-butyl ester (1.71 g, 10.0 mmol) in DCM (15 ml) was added PyBOP (4.5 g, 8.65 mmol) and DIPEA (2.24 ml, 16.5 mmol). The reaction was stirred at rt for 16 hours and then washed with aq. sodium bicarbonate, dried and evaporated. Purification by FC ($SiO_2$, ethyl acetate/MeOH) afforded the intermediate ester 2.68 g. To a stirred solution of the ester (1.3 g) dissolved in DCM (7 ml) at 0 ˚C was added tri-isopropylsilane (0.1 ml) and then TFA (10 ml). The mixture was stirred for four hours. The solvent was evaporated and the mixture was partitioned between ethyl acetate and 0.8 M aq HCl. The organics were separated, dried and evaporated to generate the product (800 mg, 61% over two steps).

$^1$H NMR (400 MHz, Chloroform-*d*) δ 9.33 (s, 0.2H), 9.05 – 8.96 (m, 0.8H), 8.37 – 8.23 (m, 2H), 8.22 – 8.14 (m, 1H), 7.94 – 7.85 (m, 1H), 7.79 (ddd, *J* = 8.5, 6.9, 1.5 Hz, 1H), 7.69 – 7.60 (m, 1H), 4.76 – 4.67 (m, 0.8H), 4.63 – 4.56 (m, 0.2H), 4.54 – 4.24 (m, 2H), 3.86 – 3.53 (m, 2H), 2.49 – 1.91 (m, 4H).

LCMS m/z 328.1 $(M+H)^+$

***N*-(2-((*S*)-2-(((*S*)-1-(4-methoxyphenyl)ethyl)carbamoyl)pyrrolidin-1-yl)-2-oxoethyl)quinoline-2-carboxamide (6)**

Methoxy-(*S*)-α-methylbenzylamine (45 µL, 0.31 mmol) was coupled to (quinoline-2-carbonyl)glycyl-*L*-proline (50 mg, 0.15 mmol) according to the general procedure A to give the product as a clear oil (39 mg, 55%).

$^1$H NMR (400 MHz, Chloroform-*d*) δ 9.01 – 8.90 (m, 1H), 8.31 (d, *J* = 8.4 Hz, 1H), 8.25 (d, *J* = 8.4 Hz, 1H), 8.19 (dq, *J* = 8.6, 0.9 Hz, 1H), 7.90 (dd, *J* = 8.1, 1.5 Hz, 1H), 7.79 (ddd, *J* = 8.4, 6.9, 1.5 Hz, 1H), 7.65 (ddd, *J* = 8.2, 6.9, 1.2 Hz, 1H), 7.28 – 7.23 (m, 2H), 7.18 (d, *J* = 7.8 Hz, 1H), 6.86 – 6.79 (m, 2H), 5.03 (p, *J* = 7.1 Hz, 1H), 4.62 (dd, *J* = 8.2, 2.2 Hz, 1H), 4.35 (t, *J* = 4.8 Hz, 2H), 3.73 (s, 4H), 3.57 (td, *J* = 9.4, 7.0 Hz, 1H), 2.45 – 2.36 (m, 1H), 2.31 – 2.16 (m, 1H), 2.12 – 2.02 (m, 1H), 1.99 – 1.87 (m, 1H), 1.49 (d, *J* = 7.0 Hz, 3H).

LCMS *m/z* 461.4 $(M+H)^+$

*N*-(2-((2*S*,4*R*)-2-(((*S*)-1-(2-chloro-4-methoxyphenyl)ethyl)carbamoyl)-4-hydroxypyrrolidin-1-yl)-2-oxoethyl)-6-fluoroquinoline-2-carboxamide CAM833A

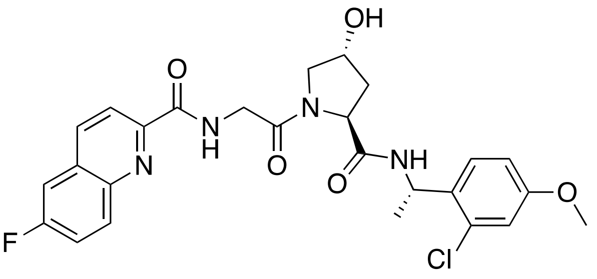

**(*S*)-1-(2-Chloro-4-methoxyphenyl)ethan-1-amine**
2-Chloro-4-methoxybenzyaldehyde (3.4 g, 19.9 mmol), (*R*)-(+)-t-butanesulfinamide (2.4 g, 19.9 mmol) were added to THF (100 mL) and $Ti(OEt)_4$ (10.0 g, 43.8 mmol) was added. The reaction was heated at reflux overnight under $N_2$, cooled to temperature and brine (100 mL) and ethyl acetate (100 mL) added. The mixture was filtered through celite, washing the celite with ethyl acetate (100 mL). The separated organic layer was dried ($Na_2SO_4$), filtered, and the solvent removed in vacuo to give the sulfinimide as a white solid (4.9 g, 90%). LCMS[b] *m/z* 274.2 $(M+H)^+$
A portion of sulfinimide (2.5 g, 9.1 mmol) was dissolved in DCM (60 mL) and cooled to -50 ˚C. MeMgBr (6.0 mL, 3.2 M in 2-methyl THF, 19.2 mmol) was added dropwise and the reaction mixture allowed to warm to rt overnight. Saturated aq. $NH_4Cl$ (50 mL) was added, stirred for 5 mins and extracted with DCM (2 x 50 mL). The combined organic layers were dried through a hydrophobic frit and purified by FC ($SiO_2$, 12-100% ethyl acetate:pet ether 40-60) to give the sulfinamine as a clear oil (2.5 g, 94%). The DR following the Grignard addition step was 98:2 as determined by $^1H$ NMR.
$^1H$ NMR (400 MHz, Chloroform-*d*) δ 7.31 (d, *J* = 8.7 Hz, 1H), 6.89 (d, *J* = 2.6 Hz, 1H), 6.80 (dd, *J* = 8.6, 2.6 Hz, 1H), 4.96 (qd, *J* = 6.7, 4.3 Hz, 1H), 3.78 (s, 3H), 1.52 (d, *J* = 6.7 Hz, 3H), 1.19 (s, 9H).
The sulfinamine (2.5 g, 8.6 mmol) was dissolved in 1,4-dioxane (60 mL) and HCl (8 mL, 4N in 1,4-dioxane) was added dropwise. After stirring for 1 hour at room temperature, the solvent was removed *in vacuo*, water (40 mL) and DCM (40 mL) were added and the organic layer discarded. The pH of the aqueous layer was adjusted to ~14 with NaOH pellets, extracted with DCM (2 x 50 mL) and the solvent removed to give the product amine as a clear oil (1.6 g, 100%).
$^1H$ NMR (400 MHz, Chloroform-*d*) δ 7.35 (d, *J* = 8.7 Hz, 1H), 6.82 (d, *J* = 2.6 Hz, 1H), 6.75 (dd, *J* = 8.6, 2.6 Hz, 1H), 4.45 – 4.38 (m, 1H), 3.72 (s, 3H), 1.30 (d, *J* = 6.6 Hz, 3H).

**(6-Fluoroquinoline-2-carbonyl)glycine**
6-Fluoroquinoline-2-carboxylic acid (7 g, 37 mmol) and glycine methyl ester hydrochloride (5.1 g, 40 mmol) were dissolved in DCM (180 mL) and DIPEA (14.8 mL, 80.5 mmol) and cooled to 0 ˚C. Over 30 minutes PyBOP (21 g, 40 mmol) was added portionwise to the solution and stirring continued for 16 h allowing the reaction to warm to rt. The yellow solution was then concentrated in vacuo and the resulting oil diluted in ethyl acetate (300 mL). The organic solution was then washed with water (4 x 100 mL), dried over $MgSO_4$ and purified by FC ($SiO_2$). The resultant methyl ester was dissolved in a MeOH, THF, water mix (1:2:1; 120 mL) and lithium hydroxide added. The solution was stirred for 1h before concentrating in vacuo and acidifying with HCl (3M aqueous). The suspension was then extracted with ethyl acetate (3 x 150 mL) and the combined organics dried and concentrated to give product (7.6 g, 84%) as a white solid.
$^1H$ NMR (400 MHz, Chloroform-*d*) δ 8.65 – 8.57 (m, 1H), 8.30 (dd, *J* = 8.5, 0.8 Hz, 1H), 8.25 (dd, *J* = 8.6, 0.8 Hz, 1H), 8.14 (ddt, *J* = 9.3, 5.5, 0.7 Hz, 1H), 7.54 (ddd, *J* = 9.2, 8.2, 2.8 Hz, 1H), 7.48 (dd, *J* = 8.7, 2.8 Hz, 1H), 4.24 (d, *J* = 5.5 Hz, 2H).
LCMS *m/z* 249.0 $(M+H)^+$

**Benzyl (2*S*,4*R*)-1-((6-fluoroquinoline-2-carbonyl)glycyl)-4-hydroxypyrrolidine-2-carboxylate**

Prepared according to general procedure B using (6-fluoroquinoline-2-carbonyl)glycine (900 mg, 3.6 mmol), benzyl (2*S*,4*R*)-4-hydroxypyrrolidine-2-carboxylate (1 g, 4.0 mmol), DIPEA (2.4 mL, 18 mmol), DCM (20 mL) and PyBOP (1.9 g, 4.0 mmol), purified by FC ($SiO_2$, 2-10% MeOH in DCM) to give the product (1.6 g, 98%) as a white solid.

$^1$H NMR (400 MHz, Chloroform-*d*) δ 8.99 – 8.85 (m, 1H), 8.31 – 8.20 (m, 2H), 8.21 – 8.12 (m, 1H), 7.54 (ddd, *J* = 9.3, 8.2, 2.8 Hz, 1H), 7.48 (dd, *J* = 8.7, 2.7 Hz, 1H), 7.43 – 7.26 (m, 5H), 5.29 – 5.13 (m, 2H), 4.76 (t, *J* = 8.0 Hz, 1H), 4.64-4.61 (m, 1H), 4.44 (d, *J* = 4.8 Hz, 0.4H), 4.39 (d, *J* = 4.8 Hz, 0.6H), 4.28 (d, *J* = 4.5 Hz, 0.6H), 4.24 (d, *J* = 4.5 Hz, 0.4H), 3.80 (d, *J* = 4.4 Hz, 0.4H), 3.78 (d, *J* = 4.3 Hz, 0.6H), 3.71 – 3.60 (m, 1H), 2.42-2.36 (m, 1H), 2.09 (ddd, *J* = 13.3, 8.0, 4.8 Hz, 2H).

**(2*S*,4*R*)-1-((6-fluoroquinoline-2-carbonyl)glycyl)-4-hydroxypyrrolidine-2-carboxylic acid**

To a solution of benzyl (2*S*,4*R*)-1-((6-fluoroquinoline-2-carbonyl)glycyl)-4-hydroxypyrrolidine-2-carboxylate (1.1g, 2.45 mmol) in MeOH (24 mL) was added palladium (10% activated on charcoal; 100 mg). The resultant mixture was hydrogenated for 16 h before filtering through celite (MeOH eluent) and concentrating in vacuo. Purification by FC ($SiO_2$; 2-20% MeOH in DCM + 1% acetic acid) gave the title compound (850 mg, 97%) as a white solid.

$^1$H NMR (400 MHz, Chloroform-*d*) δ 8.90 (q, *J* = 6.8, 5.8 Hz, 1H), 8.15 (d, *J* = 8.6 Hz, 1H), 8.08 (dd, *J* = 8.6, 0.8 Hz, 1H), 8.01 (dd, *J* = 9.3, 5.2 Hz, 1H), 7.46 – 7.33 (m, 2H), 4.56 – 4.44 (m, 1H), 4.46 – 4.37 (m, 1H), 4.30 – 4.18 (m, 1H), 4.16 – 4.05 (m, 1H), 3.69 – 3.58 (m, 1H), 3.51 – 3.41 (m, 1H), 2.25 – 2.15 (m, 1H), 2.06 – 1.94 (m, 1H), 1.36 – 1.17 (m, 1H).

LCMS *m/z* 362.2 $(M+H)^+$

***N*-(2-((2*S*,4*R*)-2-(((*S*)-1-(2-chloro-4-methoxyphenyl)ethyl)carbamoyl)-4-hydroxypyrrolidin-1-yl)-2-oxoethyl)-6-fluoroquinoline-2-carboxamide CAM833A**

(2*S*,4*R*)-1-((6-fluoroquinoline-2-carbonyl)glycyl)-4-hydroxypyrrolidine-2-carboxylic acid (3.6 g, 10.1 mmol) and (*S*)-1-(2-chloro-4-methoxyphenyl)ethan-1-amine (1.6 g, 8.61 mmol) were dissolved in DCM (50 mL), DIPEA (8.0 mL, 45.9 mmol) was added and the reaction mixture was cooled to 0 ˚C. HBTU (4.0 g. 10.6 mmol) was added portion-wise over 30 mins and the reaction mixture was allowed to warm to room temperature and stirred for 3 hours. The resulting precipitate was removed by filtration, and dissolved in DCM (200 mL) and washed with water (200 mL). Some precipitation occurred during the aqueous wash, which was collected by filtration, dissolved in ethyl acetate (150 mL) and washed with water (2 x 100 mL). The DCM organic layer was concentrated, and ethyl acetate (150 mL) was added and washed with water (2 x 100 mL). The organic layers were combined, dried with brine and magnesium sulfate and the solvent removed *in vacuo* to give the product as a white solid (2.8 g, 61%).

$^1$H NMR (500 MHz, DMSO-*d6*) δ 9.00-8.94 (m, 1H), 8.83 (d, J = 7.5 Hz, 0.3H), 8.58 (d, J = 8.5 Hz, 1H), 8.44 (d, J = 7.5 Hz, 0.7H), 8.25-8.18 (m, 2H), 7.93 (dd, J = 9.0, 2.5 Hz, 1H), 7.82 (m, 1H), 7.36 (d, J = 8.5 Hz, 0.3H), 7.29 (d, J = 8.5 Hz, 0.7H), 7.00 (d, J = 2.5 Hz, 0.3H), 6.95 (d, J = 2.5 Hz, 0.7H), 6.94 (dd, J = 2.5 Hz, 0.3H), 6.86 (dd, J = 8.5 Hz, 2.5 Hz, 0.7H), 5.19 (m, 1H), 5.08 (m, 0.7H), 4.60 (t, J = 7.5 Hz, 0.3H), 4.42 (t, J = 7.5 Hz, 0.7H), 4.35 (m, 0.7H), 4.27-4.12 (m, 2H), 3.87 (dd, 17.0, 5.5 Hz, 0.3H), 3.75 (s, 0.9H), 3.71 (s, 2.1H), 3.70-3.39 (m, 2H), 2.25 (m, 0.3H), 2.06 (m, 0.7H), 1.94 (m, 0.3H), 1.78 (m, 0.7H), 1.39 (d, J = 7.0 Hz, 0.9H), 1.30 (d, J = 7.0 Hz, 2.1H).

LCMS[b] *m/z* 529.3 $(M+H)^+$
